# Evolutionary responses to increased opportunity for sexual selection in yeast

**DOI:** 10.1101/2025.11.03.686343

**Authors:** Linnea Sandell, Anna L. Bazzicalupo, Sarah P. Otto, Nathaniel P. Sharp

## Abstract

Sexual selection contributes to biodiversity and the costs and benefits of sexual reproduction. In organisms where sex is infrequent, these impacts of sexual selection are likely to be limited. An increased frequency of obligate sex would increase the opportunity for sexual selection, which could promote the evolution of sexual traits and sexual differentiation. To study these dynamics, we conducted experimental evolution in the yeast *Saccharomyces cerevisiae*, which is predominantly asexual, with two isogamous mating types. We used selectable markers to impose frequent obligate sex in 96 populations. We manipulated the opportunity for sexual selection by imposing skewed mating-type ratios and by altering the degree to which mating occurred by selfing versus outcrossing. After just ten sexual cycles, we observed evolution in growth, cell size, pheromone production, and mating, with the mating types responding asymmetrically, but little evolutionary change in sporulation rate. Mating type dimorphism increased, with evident trade-offs between growth, attractiveness, and cell size. Genome sequences from a subset of populations revealed many mutations affecting sex-related genes. Unexpectedly, the selfing populations evolved to become sporulation-competent haploids, unlinking meiosis from ploidy change. Our results illustrate that sexual differentiation can evolve rapidly in response to an increased opportunity for sexual selection.

## Introduction

Most organisms reproduce sexually at least some of the time, which creates opportunities for sexual selection. Sexual selection is responsible for much of nature’s diversity: from the hypersensitivity of moth antennae for the pheromone of the opposite sex, to the stunning precision with which corals coordinate the release of gametes (Andersson 1994). In facultatively sexual species like the budding yeast *Saccharomyces cerevisiae*, alleles affecting sexual traits may be hidden from selection during asexual generations, thus reducing the opportunity for sexual selection. Traits subject to selection more rarely might degrade due to a lack of constraint, where mutations affecting the trait are selectively neutral, or because of pleiotropic trade-offs between sexual and non-sexual traits. Laboratory strains of *S. cerevisiae* are often maintained with little or no sex, and domesticated strains typically show a lower propensity for sporulation (meiosis) compared to wild isolates (Gerke et al. 2006). A better understanding of how alternative modes of reproduction evolve and interact might be gained by forcing *S. cerevisiae* to evolve under an increased opportunity for sexual selection.

The properties of *S. cerevisiae* have made it a popular system for evolutionary studies of reproductive mode. Natural isolates of yeast are most often diploid (Peter et al. 2018). In response to nutrient starvation, diploid cells undergo meiosis and sporulation to produce haploid offspring, which bear either *MAT*a or *MAT⍺* mating types. The two mating types look identical but produce different pheromones, called a- and *⍺*-factor, respectively. They also possess pheromone receptors for the factor produced by the opposite mating type. After sporulation, haploid spores germinate and start growing once favorable conditions return. Haploid yeast can propagate asexually but arrest growth and mate when in the presence of the opposite mating type. Because each diploid meiosis results in two spores of each mating type, it is believed to be common for siblings to mate with one another immediately upon germination, referred to as automixis (Knop 2006). If related spores are separated, for example due to passage through an insect gut (Reuter et al. 2007), haploid cells can still switch mating type during mitosis, allowing for mating and diploid formation, referred to as haplo-selfing (Hittinger 2013). Outcrossing sex (mating between lineages that are not immediate relatives) is estimated to have occurred only once every 50000 asexual generations in this species (Ruderfer et al. 2006; Magwene et al. 2011). In the laboratory, asexual haploids are maintained by eliminating the gene that allows for mating type switching, *HO*, and by maintaining *MATa* and *MAT⍺* populations separately. The opportunity for sexual selection can be manipulated by controlling the time allowed for mating and the mating-type ratio.

While sexual reproduction in *S. cerevisiae* may be relatively rare, there is evidence from laboratory studies that pheromone production is not only an indicator of mating compatibility, but a sexually selected trait (Rogers and Greig 2009; Smith et al. 2014), and that mate choice can improve population mean fitness (Strauss et al. 2024). There is also evidence that sexual traits in this species are subject to life history trade-offs. Deleting the genes that produce pheromones (*mfa1*, *mfa2*, *mfα1*, and *mfα2*) increases growth rate in both mating types (Smith and Greig 2010). Lang et al. (2009) found that sterile yeast have increased asexual growth, probably due to a decrease in costly expression of genes in the mating pathway (Lang et al. 2009). By contrast to the strong trade-offs observed between mating and growth, deleting genes responsible for meiosis did not benefit asexual growth (Goddard et al. 2005). While sporulation rate often declines in domesticated and asexually propagated strains (e.g., (Zeyl et al. 2005)), trade-offs may not be involved. Rather, mutations that interfere with meiosis and sporulation may be effectively neutral and accumulate under asexual culturing conditions (Zeyl et al. 2005). Attributes that increase competitiveness during mating in yeast also exhibit trade-offs. (Reding et al. 2013) found that populations experiencing weak mating competition adapted faster to a new environment compared to populations that underwent strong sexual selection. Likewise, impaired mating capability in yeast can be recovered only if there is strong competition for mates (Rogers and Greig 2009).

Our aim was to measure changes in growth rate, mating rate, sporulation rate, and cell size following evolution with different opportunities for sexual selection in initially isogenic populations of yeast. We manipulated the ratio of the two mating types to alter the expected intensity of sexual selection, and we imposed obligate sex by selecting for diploids immediately after mating. Previous studies of mating system in yeast have either had equal mating-type ratios (Zeyl et al. 2005) or enforced strong sexual selection for only one of the two mating types (Rogers and Greig 2009; Reding et al. 2013). In the presence of a biased mating-type ratio, we predicted that the more common mating type might evolve increased mate attraction or evolve to produce more numerous and smaller gamete-like cells than competing lineages.

We allowed yeast to evolve under alternative mating conditions, with varying levels and directions of sexual competition (Fig. 1). We used the experimental system developed by (McDonald et al. 2016) to force replicate populations to go through frequent rounds of sex with controlled periods of haploid growth, mating, diploid growth, and sporulation. During mating, *MAT*a and *MAT⍺* cells were combined either in equal ratios, or with a surplus of *MAT*a (a-biased), or with a surplus of *MAT⍺* (*⍺*-biased). We refer to these treatments as “outcrossing”. As an alternative treatment, we included populations where the mating types were not isolated during germination but rather were allowed to immediately mate within tetrads, thereby imposing selection for sex but with a lower degree of sexual competition. We refer to this treatment as “selfing”.

**Figure 1.**
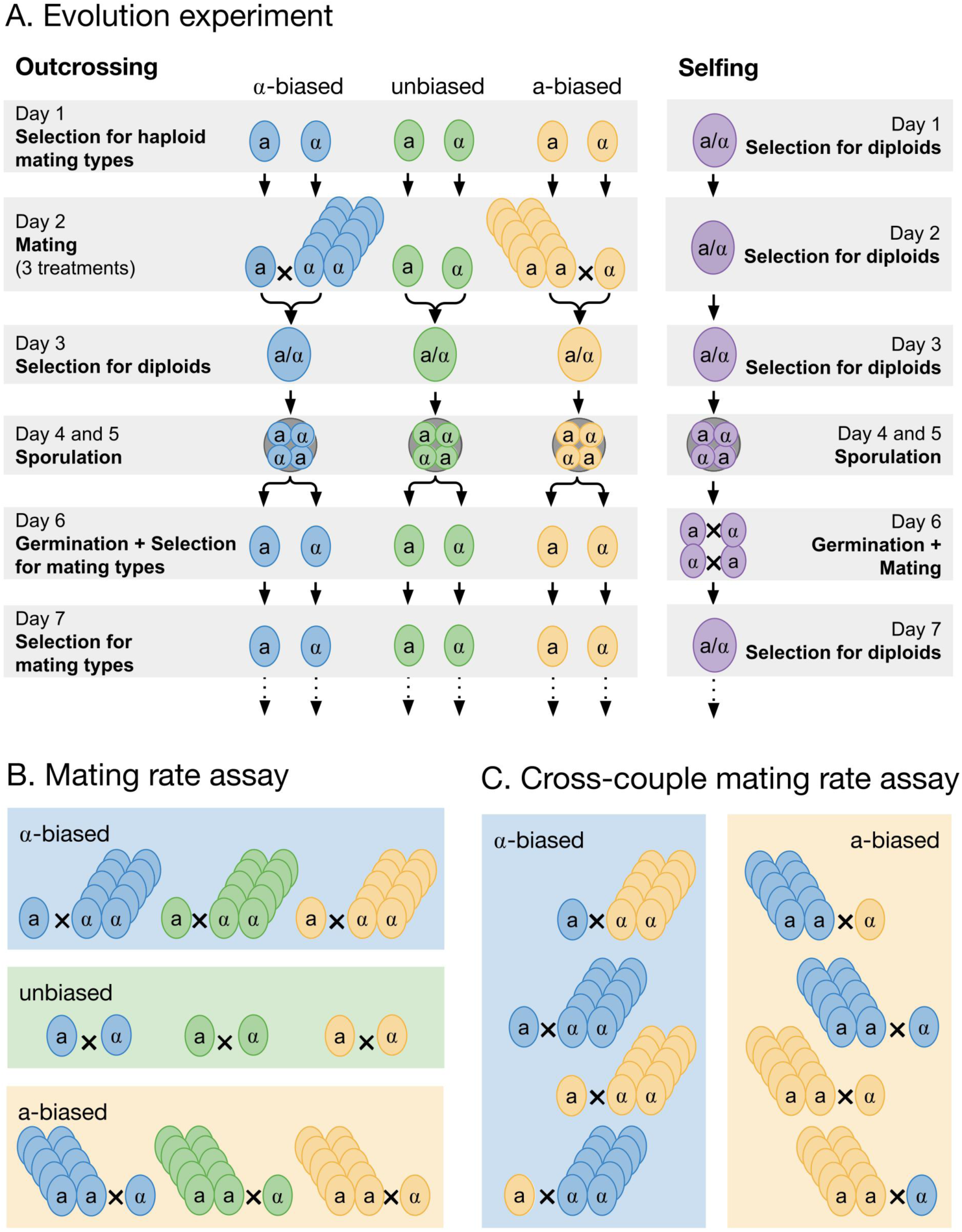
Schematic of experimental evolution and mating rate assays. (A) General protocol for maintaining outcrossing and selfing populations. Asexual growth occurred on all days apart from days 4-6. (B) General protocol for testing mating rates under alternative mating-type ratio conditions. (C) General protocol for testing mating rates with and without co-evolution of the two mating types. The background color in B and C represent the ratio of *MAT*a and *MAT*⍺ in the mating culture (blue is excess of *MAT*⍺, green is unbiased, and yellow is excess of *MAT*a). The color of the cells signifies the evolutionary history of the line used.

We predicted that mating efficiency would increase to different extents in the different treatments. Populations with skewed mating ratios were expected to obtain mutations favoring competitive mating ability in the common mating type cells, *e.g.* by increasing production of mating pheromone or sensitivity to the opposite mating factor. We also investigated whether both mating types of a population evolved to be better at mating in general, regardless of the evolutionary history of their partner, or whether the mating types coevolved to the particular mating condition.

After ten cycles of sexual reproduction, we measured the mating rate of genotypes sampled from each population under the three different mating ratios to test whether they had adapted to their mating treatment. We found that mating rate increased in all populations compared to the ancestor when mated at an equal ratio but found large differences in mating rate between treatment groups under biased mating-type ratios. We discuss the differences in the mating machinery of *MAT*a and *MAT*α cells that could drive this pattern. We also measured changes in asexual growth rate in both the diploid and haploid versions of our populations, as well as changes to cell size and pheromone production in both mating types. Finally, we sequenced genomes from a subset of populations to infer the types of genetic pathways subject to selection in the experiment. Our results illustrate that increased sexual selection and outcrossing can promote mating type differentiation and reveal trade-offs between life history traits.

## Methods

### Strains and culture conditions

We used the strain MJM64, with genotype *MAT*a *YCR043C*::KanMX, *ho*, Pr*_STE5_*-*URA3*, *ade2-1*, *his3*Δ::3xHA, *leu2*Δ::3xHA, *trp1*-1, can1::Pr*_STE2_*-*HIS3*-Pr*_STE3_*-*LEU2* and strain MJM36 with genotype *MAT⍺ YCR043C*::HphMX, *ho*, Pr*_STE5_*-*URA3*, *ade2*-1, *his3*Δ::3xHA, *leu2*Δ::3xHA, *trp1*-1, *can1*::Pr*_STE2_*-*HIS3*-Pr*_STE3_*-*LEU2* described in (McDonald et al. 2016), which are originally from a common W303 genetic background. These strains have been genetically modified such that several amino acid biosynthesis genes are under the control of promoters specific to certain cell types (S1 Text). The *STE5* promoter is haploid-specific, allowing haploid selection using uracil drop-out medium and counter-selection using 5-FoA; the *STE2* promoter is *MAT*a-specific, allowing selection for this mating type on histidine drop-out medium; the *STE3* promoter is MAT⍺-specific, allowing selection for this mating type on leucine drop-out medium. Additionally, the antibiotic markers KanMX and HphMX are tightly linked to *MAT*a and *MAT*⍺, respectively, and so only MATa/MAT⍺ cells can grow when both antibiotics are present. These strains allowed us to enforce an alternation of generations (McDonald et al. 2016), unlike other yeast strains available (see Fig. S1 and S1 Text). Unless otherwise noted, we incubated all experimental cultures at 30° C.

We combined selection on the auxotrophic markers and antibiotics to select for haploid cells, as detailed in Table 1. Because ammonium sulphate (AS), which is commonly used as a nitrogen source in synthetic medium, would interfere with antibiotic selection, we replaced it with monosodium glutamate (MSG) in our media (referred to as SE). Similarly, we used media containing 5-FoA and two antibiotics to select for diploids (Table 1).

**Table 1.** Media used in this study. See S1 Text for additional details.

| Medium | Selects for |
| --- | --- |
| SE<sup>a</sup> + drop-out mix lacking Ura and His + G418<sup>b</sup> | Haploid *MATa* |
| SE<sup>a</sup> + drop-out mix lacking Ura and Leu + hygromycin<sup>c</sup> | Haploid *MATα* |
| SE<sup>a</sup> + 5-FoA + G418<sup>b</sup> + hygromycin<sup>c</sup> | Diploid *MATa* / *MATα* |
| SE<sup>a</sup> + 1 % potassium acetate | Presporulation |
| Essential amino acids<sup>d</sup> + 1 % potassium acetate | Sporulation |

<sup>a</sup> Dextrose (42 g/L) + yeast nitrogen base (3.9 g/L) + monosodium glutamate monohydrate (3 g/L).
<sup>b</sup> 250 µg/mL.
<sup>c</sup> 200 µg/mL.
<sup>d</sup> 5 mg/L Ade, Ura, His, and Trp; 25 mg/L of Leu.

### Evolution experiment

We grew experimental cultures in 1 mL of liquid media in 96-well plates with shaking. We initiated 96 *MAT*a and 96 *MAT*α populations by adding 10 μL of saturated culture of each ancestor to 1 mL of medium selecting for that mating type (Table 1). Each cycle of sexual reproduction consisted of mating, sporulation, and germination with haploid growth, as follows.

#### Mating

We mated cultures of opposite mating types in each well of a 96-well plate, manipulating the mating-type ratio by altering the relative volumes of *MAT*a to *MAT⍺* culture used: either combined in an equal ratio or 1:10 ratio in favor of either *MAT*a or *MAT⍺*, with 24 replicate populations for each of these three treatments. We chose to hold constant the total number of potential diploids by diluting the rarer mating type by a constant amount, diluting the more common mating type less in the skewed mating ratio treatments to allow for more competition (Table 2; S2 Text). Although our main goal was to compare evolution under alternative mating-type ratios, we also maintained an additional 24 populations where we did not separately culture *MAT*a and *MAT⍺*, allowing self-fertilization within tetrads and their descendants (“selfing” treatment). To match the outcrossing treatments, we divided each selfing population into two wells during the periods where the outcrossing populations were grown as separate mating types and subsequently combined these subpopulations in equal ratio (similar to the equal mating ratio treatment).

**Table 2.** Mating ratios during the experiment and in post-experiment assays.

| Ratio | *MATa* [mL]<sup>a</sup> | *MATα* [mL]<sup>a</sup> | *MATa* dil<sup>b</sup> | *MATα* dil<sup>b</sup> | Cell dil<sup>b</sup> |
| --- | --- | --- | --- | --- | --- |
| a-biased | 1 | 0.1 | 5.5 | 55 | 5 |
| unbiased | 0.1 | 0.1 | 50 | 50 | 25 |
| α-biased | 0.1 | 1 | 55 | 5.5 | 5 |

<sup>a</sup> Volumes sampled from the saturated cultures of haploids grown in SE.
<sup>b</sup> Final dilutions of each mating type or of the total number of cells (see details on dilution series in S2 Text).

**Table 3.**
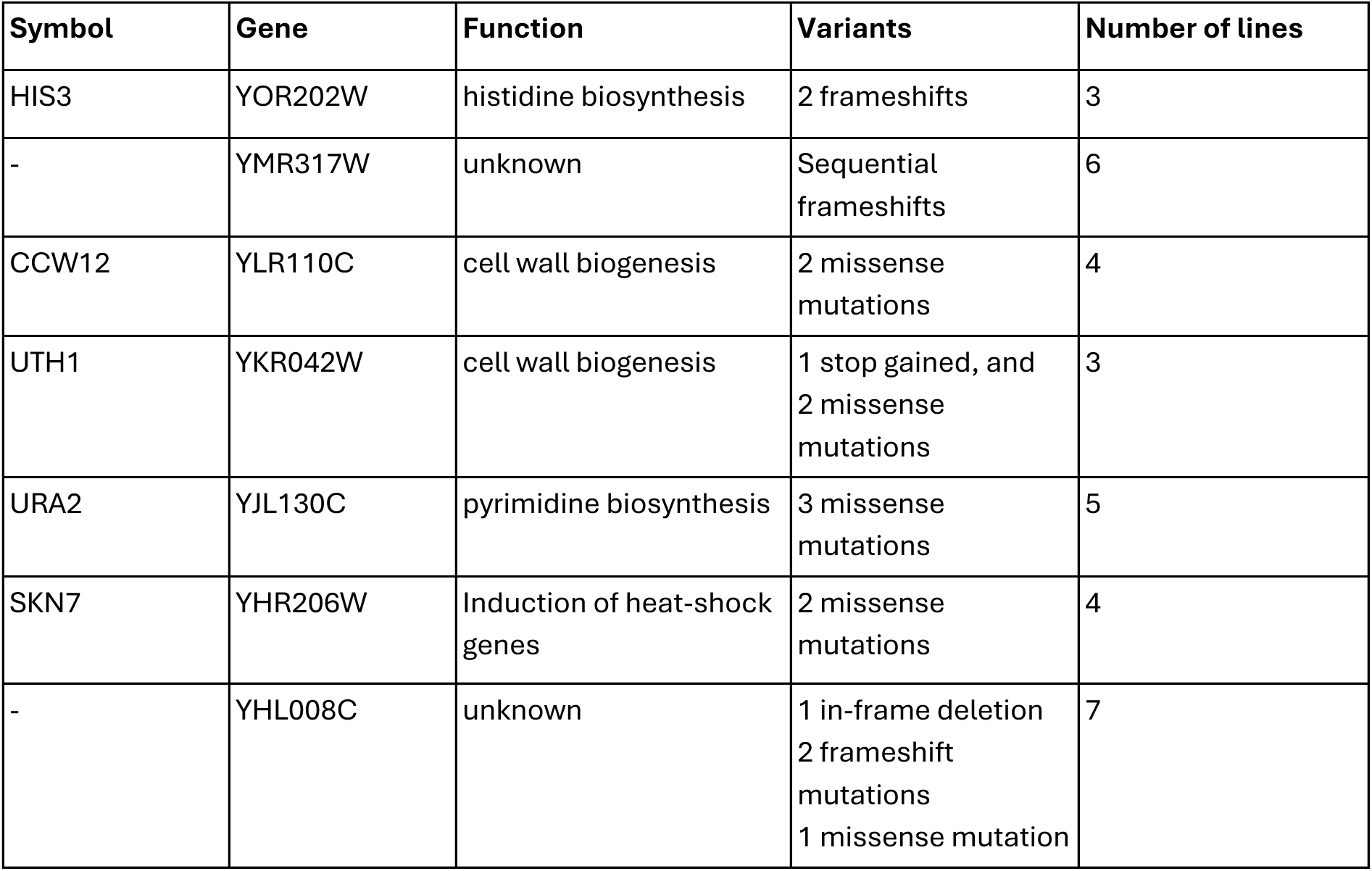
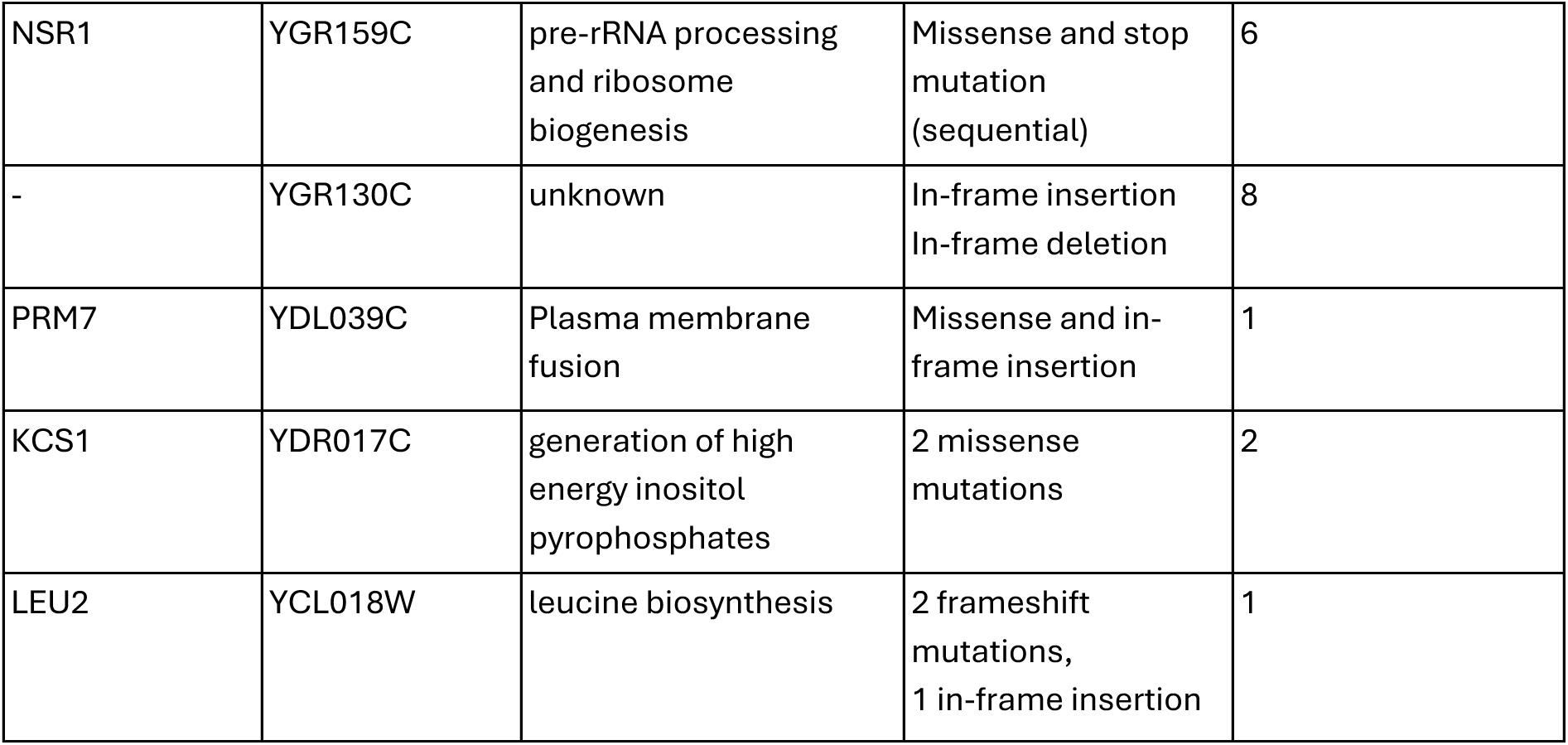
Genes with multiple hits in the 12 outcrossing samples sequenced.

| Symbol | Gene | Function | Variants | Number of lines |
| --- | --- | --- | --- | --- |
| HIS3 | YOR202W | histidine biosynthesis | 2 frameshifts | 3 |
| - | YMR317W | unknown | Sequential frameshifts | 6 |
| CCW12 | YLR110C | cell wall biogenesis | 2 missense mutations | 4 |
| UTH1 | YKR042W | cell wall biogenesis | 1 stop gained, and 2 missense mutations | 3 |
| URA2 | YJL130C | pyrimidine biosynthesis | 3 missense mutations | 5 |
| SKN7 | YHR206W | Induction of heat-shock genes | 2 missense mutations | 4 |
| - | YHL008C | unknown | 1 in-frame deletion<br>2 frameshift mutations<br>1 missense mutation | 7 |
| NSR1 | YGR159C | pre-rRNA processing and ribosome biogenesis | Missense and stop mutation (sequential) | 6 |
| - | YGR130C | unknown | In-frame insertion<br>In-frame deletion | 8 |
| PRM7 | YDL039C | Plasma membrane fusion | Missense and in-frame insertion | 1 |
| KCS1 | YDR017C | generation of high energy inositol pyrophosphates | 2 missense mutations | 2 |
| LEU2 | YCL018W | leucine biosynthesis | 2 frameshift mutations,<br>1 in-frame insertion | 1 |

#### Sporulation

We selected for diploids for one day using diploid-selective media (Table 1), which should also eliminate any remaining haploid cells. Following diploid selection, we incubated cells in pre-sporulation medium (Table 1) for 15 hours; we then transferred cultures to individual 10 mL plastic tubes, centrifuged in order to remove the pre-sporulation medium, and replaced it with 1 mL of sporulation medium (Table 1). We incubated these cultures in a roller drum at room temperature for 2 d.

#### Germination

We transferred the sporulated cultures to microcentrifuge tubes, centrifuged, and removed supernatant. To select against unsporulated cells, we treated the sporulated cultures with 5 μL of 5 U/μL of zymolyase in 45 μL of sterile water (giving 0.5 U/μL) for 20 minutes at 35° C and then put the tubes on ice. To disrupt tetrads in the outcrossing treatments, we sonicated the cultures using a Bioruptor machine with 30 s sonication followed by 30 s rest, repeated ten times. We then divided each sonicated culture, with 20 μL used to inoculate 1 mL of *MAT*a-specific medium and 20 μL used to inoculate 1 mL of *MAT⍺*-specific medium. We did not sonicate the selfing populations after the zymolyase digestion, as we wanted the tetrads to remain intact to allow for within-tetrad mating. Rather, we split the culture into two wells as noted above, transferring 20 μL culture into 1 mL of non-selective SE medium. After 2 d of germination and growth in mating type-specific media for the outcrossing populations or 1 d of germination and growth in SE medium followed by 1 d of growth in diploid-specific medium for the selfing populations (days 6 and 7 in Fig. 1A), we diluted all populations 1:100 into SE medium and started the next cycle of sexual reproduction.

Using the procedure described above, we subjected the experimental populations to ten cycles of sexual reproduction, each of which took 7 days to complete (Fig. 1). We stored and froze samples from these populations during each cycle at day two before mating (haploids) and day 3 before transfer to presporulation medium (diploids) and preserved all cultures after the seventh day of the tenth sexual cycle. Given a doubling time of 90 minutes under optimal conditions, we expect that as many as 64 generations of asexual growth occurred per experimental cycle (7 days). Although population sizes in many of these stages of growth were likely large (yeast cultures typically grow to ∼3 × 10^8^ cells/mL), the effective population size during the experiment would have been much smaller, due to bottlenecks during mating and sporulation. While the effective population size was not measured, the evolutionary and genomic responses documented below indicate that sufficient genetic variation arose during the experiment to allow a response to selection.

We compare results between the outcrossing and selfing populations, and among the outcrossing treatments, all with reference to the properties of their ancestors from frozen stock. We did not compare the outcrossing lines to an asexual control because the experimental design required passage through sporulation media and enforced an alternation of haploid and diploid generations through selective media (Table 1). Asexual propagation would thus have required different media (or a complex set of genetic manipulations allowing growth without sex in all media), confounding differences in mode of reproduction. We were also concerned about confounding effects of spontaneous ploidy changes when propagating yeast asexually, as observed in previous experiments (Gerstein et al. 2006; Gerstein and Berman 2015; Fisher et al. 2018; Harari et al. 2018; Gerstein and Sharp 2021).

### Preservation of evolved and ancestral strains

At the end of the experiment, we streaked frozen cultures from day 3 (diploids) and or day 7 (haploids) of cycle 10 on SE plates and used single colonies to inoculate 2 mL of SE medium to obtain isogenic representatives for each population. After 2 d of growth, we combined each culture with glycerol and froze at –80° C. We also performed this procedure for eight independently grown cultures of the ancestral haploid *MAT*a, the ancestral haploid *MAT*α, and the ancestral diploid formed by mating the two haploid ancestors. We used the ancestral diploid strain as a control in flow cytometry when assaying genome size and cell size in the evolved populations. We also used the cultures of haploid ancestors in flow cytometry and in other assays of the evolved haploids. We used single colonies derived from the diploid populations of cycle 1 (day 3) to grow and freeze ancestral controls for comparison with the evolved diploid. We found evidence of contamination in one *MAT⍺* strain, which was excluded from all analyses.

### Mating rate assays

We measured mating rates under unbiased, *⍺*-biased, and a-biased ratios (Table 2), with three replicates each, after first growing haploids in SE medium overnight. After 24 h, we diluted and plated each mating culture on both permissive and diploid-selective media. The assay was conducted in 9 blocks; in the first two blocks we used SE and SE + 5-FOA + G418 + hygromycin for permissive and diploid-selective media, respectively, but replaced SE with YPAD (Yeast Peptone Dextrose plus 40 mg/L adenine) in the remaining blocks, for practical reasons (block was used as a random effect in our model). Because only a small fraction of cells were diploid, we made separate dilutions for plating on permissive and diploid-selective plates. We counted colonies on each plate after 2 d.

To calculate mating rate, we first converted colony counts into cell concentrations in the mating culture. The colony count on diploid-selective plates gives the concentration of diploid cells, D_mated_, whereas the permissive plates give the total concentration of cells, C_mated_ (a mix of haploids and diploids). The maximum concentration of diploids, D_max_, is different for the biased and unbiased mating scenarios. In the unbiased mating ratio, the final culture could (theoretically) consist of 100% diploids if all cells were mated (D_max_ = C_mated_). In contrast, due to the excess of one mating type, the maximum fraction of diploids in the biased ratio would be 10% because 10:1 mating ratio would produce at most 9:1 haploids:diploids if all the rare types mated (D_max_ = 0.1 C_mated_). We therefore calculated mating rate by dividing D_mated_ by D_max_. This calculation assumes that all cell types grew at the same rate during the 24 h mating period. Because the presence of pheromones inhibits haploid growth, diploid cells may have grown faster once produced, which would bias the estimated mating rate upwards. To avoid cases where not enough cells were plated to obtain accurate estimates, we removed plates with fewer than 500 colonies expected on the diploid-selective plate had there been 100% mating (based on D_max_/the dilution factor used for the selective plate).

Because of the way we infer mating rate (the concentration of diploids from one plate and the maximum possible concentration of diploids from another), the error distribution is not binomial (which would normally be used for sampled proportions). We thus fit a linear-mixed effect model to our data, using population ID and block as random effects and evolutionary treatment and assay mating ratio as fixed effects, allowing interactions. To disentangle the interactive effects of evolutionary treatment and assay ratio we used the *emmeans* package (Lenth 2017) to make conditional pairwise comparisons between assay ratios and treatment groups. We also tested whether changes in mating rate were due to evolutionary changes in the common or in the rare mating type by conducting mating assays with partners from different populations, considering only the biased mating-ratio treatments (S3 Text).

To estimate the mating rate in the selfing treatment, we sporulated the ancestral diploid in 16 replicates, treated the cultures with zymolyase with the same procedure as we used for the selfing treatment during the evolution experiment, and germinated them in the same ratio of treated culture to medium as we used during evolution. After 24 h of growth, we diluted the culture and spread it on permissive and diploid-selective plates. Using colony counts on the two types of plates, we calculated the mating rate as described above.

### Pheromone production assays

We quantified the production of mating pheromones on plates by placing a spot of each strain of interest on a lawn of cells from a tester strain of the opposite mating type. The tester strains, *MATa bar1* and *MAT⍺ sst2*, arrest their growth upon detection of mating pheromone from the opposite mating type, creating a “halo” of growth inhibition on the lawn surrounding the query strain. This approach is used qualitatively to determine mating type, *i.e.*, the presence of a halo indicates the query is the opposite mating type from that of the lawn strain; in this case we applied the assay quantitatively, using the radius of the halo as a measure of pheromone production.

We grew replicate populations of the query and tester strains in liquid YPAD in deep-well plates for 2 days. To prepare a tester plate (YPAD media), we combined 50 µL of a given tester strain with 50 µL of sorbitol and spread this solution on a plate using sterile glass beads. To improve the visibility of halos when the query strains were *MATa*, which produce hydrophobic pheromone, we included 0.05% Triton-X100 in the media for these test plates. We incubated test plates with lawns of the tester strains for 30-60 min before adding spots of randomly selected opposite-type query strains. After 2 d we imaged plates from below using a flatbed scanner. Using ImageJ (Fiji) (Schindelin et al. 2012), we used the oval tool to determine the area (arbitrary units) of each halo and each spot of query yeast. To calculate the width of the zone of inhibition, we used the halo area to calculate a halo radius, used the spot area to calculate a spot radius, and subtracted the spot radius from the halo radius. Using the size of the plate (which has a known diameter) from the images, we converted halo radius measurements into units of mm.

We used the *lmer* package to fit linear mixed models with halo radius as the response variable and random effects of query strain and plate, with a fixed effect of evolutionary treatment (unbiased, a-biased, *⍺*-biased, or ancestor) or evolution status (evolved vs. ancestral). We also examined correlations between halo size and mating type for *MATa* and *MAT⍺* strains from the same population.

### Sporulation rate assay and spore staining

We measured the sporulation rate of a subset of populations during the experiment (eight from the selfing treatment and eight from each of the three outcrossing treatments), taking estimates from the same set of populations at each cycle. For each sexual cycle, we sampled 10 μL of sporulated culture and counted the number of cells and the number of tetrads with the use of a hemocytometer (tetrads were counted as one cell). We counted 150.2 cells per population per assay on average (SD = 52) and estimated sporulation rate as the number of tetrads divided by the total number of cells. We fit a generalized linear model of sporulation rate on evolutionary treatment, the number of sexual cycles, and their interaction.

Following experimental evolution, we revived the diploid populations from the outcrossing treatments of both sexual cycle 1 and sexual cycle 10 from frozen samples and allowed for two days of sporulation using the protocol described above. We scored sporulation rates in six blocks, counting 302 cells per population on average (SD: 127). We found that block and experimenter explained little variance, so we excluded these random factors and fit a generalized linear model of sporulation rate as a function of evolutionary treatment or evolution status (ancestor versus evolved).

Given the evolved characteristics of the selfing populations (see Results), we considered whether these particular strains might be forming spores in a non-tetrad format that would be difficult to score in the assays described above. We applied a published spore staining protocol (Evans et al. 1949), with modifications (S4 Text), which allows spores to be visually distinguished from vegetative cells. We revived the 24 frozen selfing populations from day 7 of cycle 10, as well as 4 haploid populations and 4 diploid populations from the outcrossing treatments, following the maintenance protocols described above. We used these cultures to inoculate sporulation medium, or SE medium for comparison, incubated for 2 d (for a subset of four selfing populations that exhibited almost no tetrads the time in sporulation medium was extended to 4 d), and then performed the staining protocol (S4 Text).

### Growth rate assays

We initiated growth rate assays by inoculating 1 mL of SE with 20 μL of frozen culture stock in deep-well plates and incubating for 2 d with shaking. We then transferred these cultures to microcentrifuge tubes and held them at 4 C to arrest growth during assay preparation. For each growth rate assay, we diluted these cultures 1:121 and measured OD_600_ every 15 min in a Bioscreen C for 19-24 h with continuous shaking. We estimated growth rate as the maximum slope of a spline fit to log-transformed OD_600_ readings using the *loess* function in *R* (Gerstein 2012). We used ANOVA to test for differences in maximum growth rate among evolved populations and used Welch’s t-test to compare evolved and ancestral populations (sexual cycle 10 and 0, respectively).

### Flow cytometry

We used flow cytometry to assess cell size and genome size in putatively haploid samples of the evolved outcrossing populations and their ancestors, as well as the selfing populations and an ancestral diploid control. We prepared cells for flow cytometry by fixing with ethanol and staining with Sytox Green (Gerstein et al. 2006) (S5 Text). We ran 160 μL of each sample on an Attune NxT analyzer and used the *flowCore* package (Hahne et al. 2009) for analysis in *R*. After removing events with very low and high fluorescence or forward scatter, we took the location of the first fluorescence peak as the genome content of G1 cells. We used the average forward scatter of cells with fluorescence within 0.5 SD of the first peak as an estimate of cell size in G1 cells. We excluded two populations from this analysis due to experimental error.

### PCR of MAT locus

To determine the mating type at the *MAT* locus we performed colony PCR with forward primers specific to *MAT*a (AAGTTGCAAAGAAATGTGGC) or *MAT⍺* (AAAATGCAGCACGGAATATG) and a common reverse primer (AACAAATTGTGAAGCCGAAG).

### Mating phenotype assays

We assayed ploidy and mating type of the evolved populations by spotting 10 μL of saturated culture onto agar plates with the two mating type-specific and the diploid-specific media (S6 Text). After two days we noted whether each spot had grown or not. We performed additional mating type assays at the end of the experiment by combining *MAT⍺* cells with *MAT*a tester strains bearing *his1* and replica plating onto dropout plates lacking histidine. Because the *his3* auxotrophy in our experimental strains complements the *his1* auxotrophy in the tester strain, only diploids formed by mating can grow on this drop-out medium. These assays were not performed for MATa cells because our *MAT⍺* tester was likely contaminated (see S6 Text).

### Trait correlations and dimorphism

To explore correlations between traits (mating rate, pheromone production, cell size, and growth rate), we considered only the populations from the outcrossing treatments. We first determined mean mating rate in each assay condition, accounting for random effects by finding predicted values from a mixed-effect model and focused on the mating rate in the mating ratio under which each had evolved. We calculated the Spearman rank correlation (*r*_S_) for a pair of traits for each treatment group separately and then averaged these correlations across parallel tests to increase power. We tested for statistical significance by randomizing the association between trait values within each group to obtain a null distribution for the average rank correlation (minimum 5000 simulations per test). We quantified mating type dimorphism for pheromone production and cell size in each evolved and ancestral population by standardizing the absolute mating type difference by the mean: 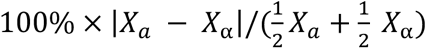, where *X* is the trait value for a given mating type; we compared this metric among groups using linear models.

### Genome analyses

To gain insight into the molecular basis of evolved phenotypes we conducted whole genome sequencing of 15 total strains including three pairs of populations from each of the biased mating type treatments (both the MATa and MAT*⍺* strain of the same evolved population, 12 strains in total), two populations from the selfing treatment, as well as the ancestral diploid strain. (We later discovered that one of the putatively MAT⍺ strains sequenced was actually a selfing strain, due to an error.) We revived frozen cultures, grew them in liquid YPAD for 48 h, and extracted DNA (Sambrook and Russell 2006). The libraries were prepared with an Illumina DNA Prep kit; some were sequenced with NextSeq paired-end 75 bp reads and others with MiSeq paired-end 150 bp reads at the Sequencing and Bioinformatics Consortium (University of British Columbia, Vancouver, Canada).

We processed the sequence data by applying cutadapt 4.6 with Python 3.9.18 (Martin 2011) to Illumina adapter sequences, mapped trimmed sequences to the S288C reference genome R64 (RefSeq assembly GCF_000146045.2, which is MAT*⍺*), using BWA mem (bwa-0.7.17-r1188; (Li and Durbin 2009)), sorted and indexed files using samtools 1.19 (Li et al. 2009), removed duplicate reads using MarkDuplicates from the Genome Analysis Toolkit (GATK) v4.4.0.0 (DePristo et al. 2011), Picard version 3.0.0, and analyzed coverage using bedtools *genomecov* v2.31.1 (Quinlan and Hall 2010). To call variants we used GATK HaplotypeCaller (Poplin et al. 2018) with ploidy set to 1, followed by GenotypeGVCF. To analyze the variants we transformed the gvcf into a table using GATK VariantsToTable. To check for variants in the MATa locus we repeated alignments to a modified version of chrIII, replacing the MAT*⍺* sequence (positions 198671–201178) with the MATa sequence. We did not identify high-quality variants in either version of the MAT locus. We retained only biallelic variants with QD > 18, FS < 10, SOR < 4, MQ > 59, –5 < MQRankSum < 5, –7.5 < ReadPosRankSum < 7.5, and QUAL > 30. We excluded putative variants where all samples either had low coverage (depth < 5) or shared the same variant, which most likely represent alleles present in the ancestral strain. We used Variant Effect Predictor to determine the genic consequences of variants and identified gene functions based on the *Saccharomyces* Genome Database (yeastgenome.org). We examined coverage for each sample and chromosome with the goal of identifying segmental duplications unique to a strain. We first calculated relative coverage values by dividing by the chromosome-wide median for that sample and then further standardized using the average relative coverage value across samples. We examined standardized coverage profiles for regions with 2-fold elevated coverage as an indication of duplication.

## Results

### Marker and mating phenotypes

Following ten sexual cycles, the haploids isolated from the outcrossing lines (day 7 in Fig. 1A) retained their expected growth properties, with the *MAT*a and *MAT⍺* cells of each treatment growing on their mating type-specific medium but not on that of the other. We also confirmed that the haploids were of the expected mating type in crosses to tester lines. By contrast, yeast from the selfing populations appeared to be mated already at this stage, exhibited the same growth pattern on the selective plates as our control diploid, and did not mate with tester strains of either mating type.

### Growth rates

There were no significant differences in growth rate among the evolved populations of different treatment groups, either among the haploid or the diploid lines, in either permissive or selective media (Fig. 2). While the evolved populations grew similarly across mating-type ratios, they often grew faster than their ancestors, particularly in the selective media. The evolved *MAT*a cells exhibited a higher growth rate than the ancestral strain in the mating-type selective medium, but not the permissive medium (*t* = 6.69, *P* < 10^−5^ and *t* = 0.62, *P* = 0.54, respectively; Fig. 2A). The evolved *MAT⍺* cells exhibited a higher growth rate compared to the ancestor in both media (*t* = 18.63, *P* < 10^−5^ and *t* = 3.87, *P* < 0.005, respectively; Fig. 2B). The evolved diploid cells exhibited a higher growth rate compared to the ancestor in the diploid-selective medium, but not permissive medium (*t* = 4.23, *P* < 10^−3^ and *t* = 0.14, *P* = 0.89, respectively; Fig. 2C), and the same pattern was evident for the selfing populations when compared to the diploid ancestor (*t* = 5.40, *P* < 10^−5^ and *t* = 0.73, *P* = 0.47, respectively; Fig. 2C). In summary, growth rates increased during experimental evolution for all cell types when grown in the medium that was used to select for each cell type.

**Figure 2.**
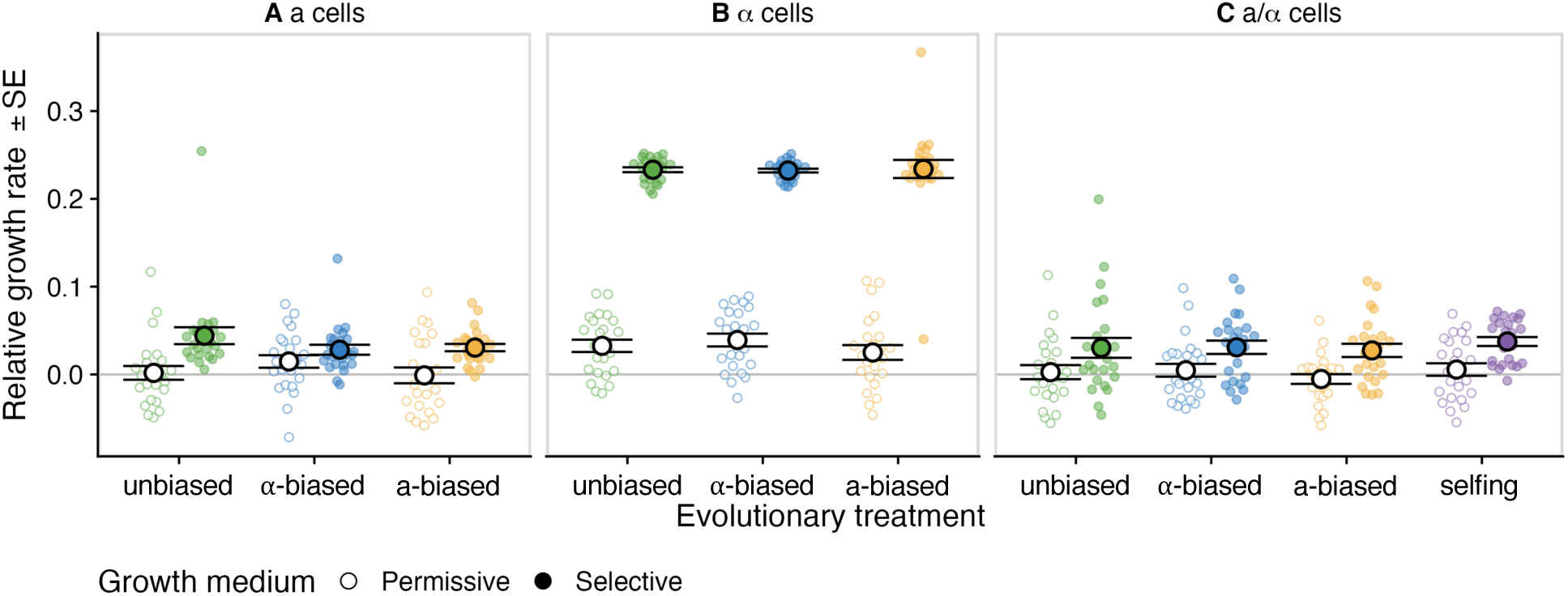
Growth rates of evolved strains in permissive and selective media. Growth rate of evolved *MAT*a cells (A), *MAT*⍺ cells (B), and *MAT*a/⍺ cells (C) is shown in permissive medium (open circles) or selective medium (*e.g.*, *MAT*a haploid selective medium for *MAT*a cells in left panel, filled circles). Black circles and error bars represent group means and standard errors.

### Mating rates

There was a significant interaction in mating rates between the evolutionary treatment (mating-type ratio) and the assay condition (*χ*^2^ = 40.128, *P* < 10^−5^; Fig. 3). Using the Tukey method for *P*-value adjustment, we then considered each contrast separately. We found no significant differences in the mating rate of the ancestor across the three assay conditions (all *P* > 0.05). Populations evolved in the *⍺*-biased treatment exhibited higher mating rates when tested in the *⍺*-biased assay, but no significant effect of assay condition was found for populations evolved in the a-biased treatment (*t*-ratio to unbiased: 2.51, *P* = 0.034; *t*-ratio to a-biased: 8.25, *P* < 10^−3^).

**Figure 3.**
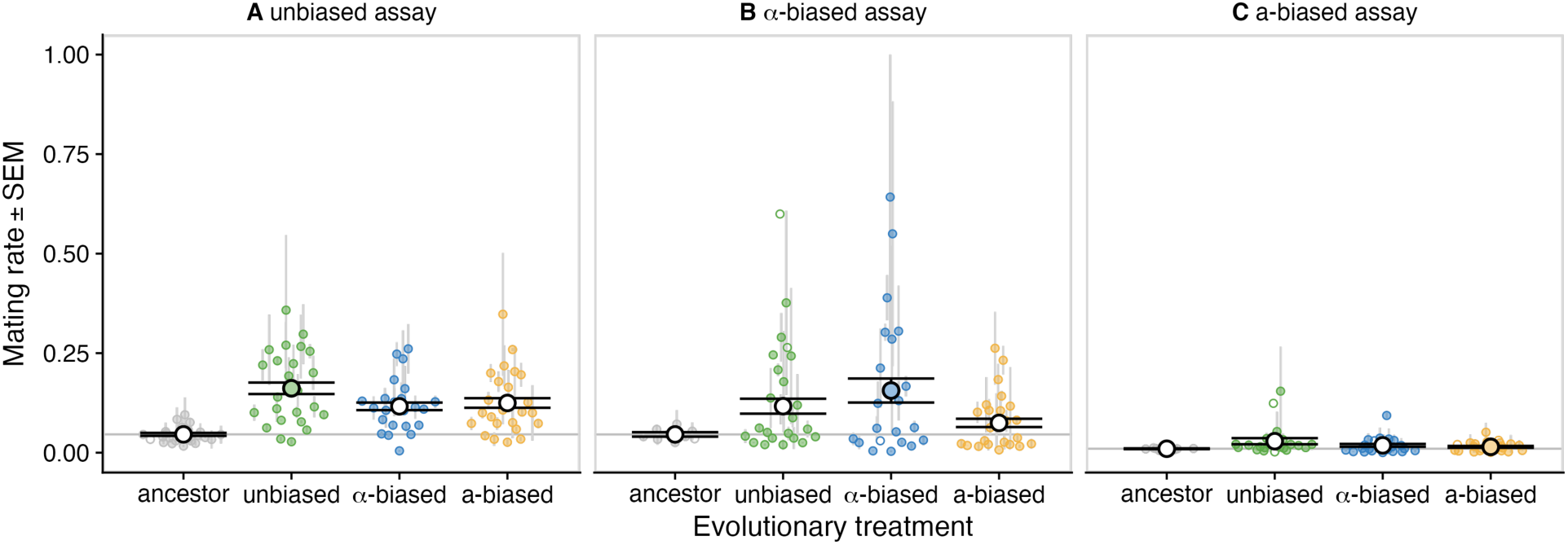
Mating rates of evolved strains. Mating rates were tested when the mating-type ratio was unbiased (A), ⍺-biased (B), or a-biased (C). Cases where the mating rate of a line was measured only once are represented by open circles; when measured more than once, solid points represent means and the grey lines represent standard errors. Black circles and error bars show group means and standard errors, with the circle filled for the treatment group that evolved under the assayed condition.

When mated in an unbiased ratio, the evolved populations of all treatments outperformed the ancestor but did not differ from one another (Fig. 3A). While the ancestor showed a mating rate of 3.9 ± 1.9% (mean ± SE), the strains evolved under unbiased, a-biased and *⍺*-biased conditions had 3-4 fold higher mating rates of 15.8 ± 1.8% (*t*-ratio: 6.64), 12 ± 1.9% (*t*-ratio: 4.52), and 11.2 ± 1.9% (*t*-ratio: 4.02), respectively (*P* < 0.005 for all comparisons). When tested in an *⍺*-biased ratio, populations that had evolved under that condition showed a higher mating rate than populations from the a-biased treatment (t-ratio: 4.02, *P* < 10^−3^; Fig. 3B). When tested in an a-biased ratio, no evolved treatment had a significantly higher mating rate than the ancestor or any other treatment group (Fig. 3C). In summary, we find that mating rates increased in all treatment groups, except under a-biased assay conditions, demonstrating that the mating rate of yeast can improve rapidly under conditions that select for more frequent sexual reproduction.

We next performed reciprocal crosses using MATa vs. MAT*⍺* partners from different evolutionary treatments (S3 Text; Fig. S2). Similar to the results above, the mating rate under the a-biased assay condition was significantly lower than in the *⍺*-biased assay condition (t = –9.16, *P* < 10^−5^). Conditional pairwise tests on the fixed effects of the model revealed that this effect was entirely driven by matings where *MAT⍺* had evolved as the common partner, which had a mean mating rate of 36.1 ± 3.3% in the *⍺*-biased ratio and 2.1 ± 3.1% in the a-biased ratio. Regardless of the assay ratio, tests where the *MAT⍺* cells came from the a-biased treatment had significantly reduced mating rate (t = –7.18, *P* < 10^− 5^), with means of 5.1 ± 2.7% in the *⍺*-biased ratio and 2.3 ± 2.5% in the a-biased ratio. We also found that *MAT⍺* cells from the *⍺*-biased treatment had a higher mating rate in its evolved condition if its mating partner also came from the same treatment (t-ratio *⍺*-biased – a-biased = 3.5, *P* = 0.011). For the *MAT*a cells from the *⍺*-biased treatment the effect was even stronger (t-ratio *⍺*-biased partner – a-biased partner = 7.18, *P* < 0.001). This did not appear to be a coevolutionary effect specific to mating pairs from the same replicate, as a likelihood ratio test excluded any significant effect of whether partners came from the same versus different evolved populations (*χ*^2^ = 0.0445, *P* = 0.83).

### Pheromone production

Pheromone production, quantified by the radius of growth inhibition on lawns of pheromone-sensitive strains (“halos,” Fig. 4), varied significantly among evolution treatments (treated as a fixed effect of unbiased, a-biased, α-biased, or ancestor) for both *MAT*a and *MAT⍺* haploid lines (*MAT*a, *P* < 0.001; *MAT⍺*, *P* < 1 × 10^−10^). Separate analyses of each of the four treatment groups and two mating types showed significant genetic variance among populations in six out of eight cases, with the exception of the ancestral *MAT⍺* strain and *MAT*a cells from the *⍺*-biased treatment. Overall, there was good support for the idea that halo sizes reflect heritable genetic variation in pheromone production. The MAT*⍺* cells appeared to show greater genetic variance (Fig. 4); averaging across the outcrossing treatments, the coefficient of genetic variance was 4.02-fold higher for *MAT⍺* strain than *MAT*a cells (significantly greater than one; bootstrap *P* = 4 × 10^−4^).

**Figure 4.**
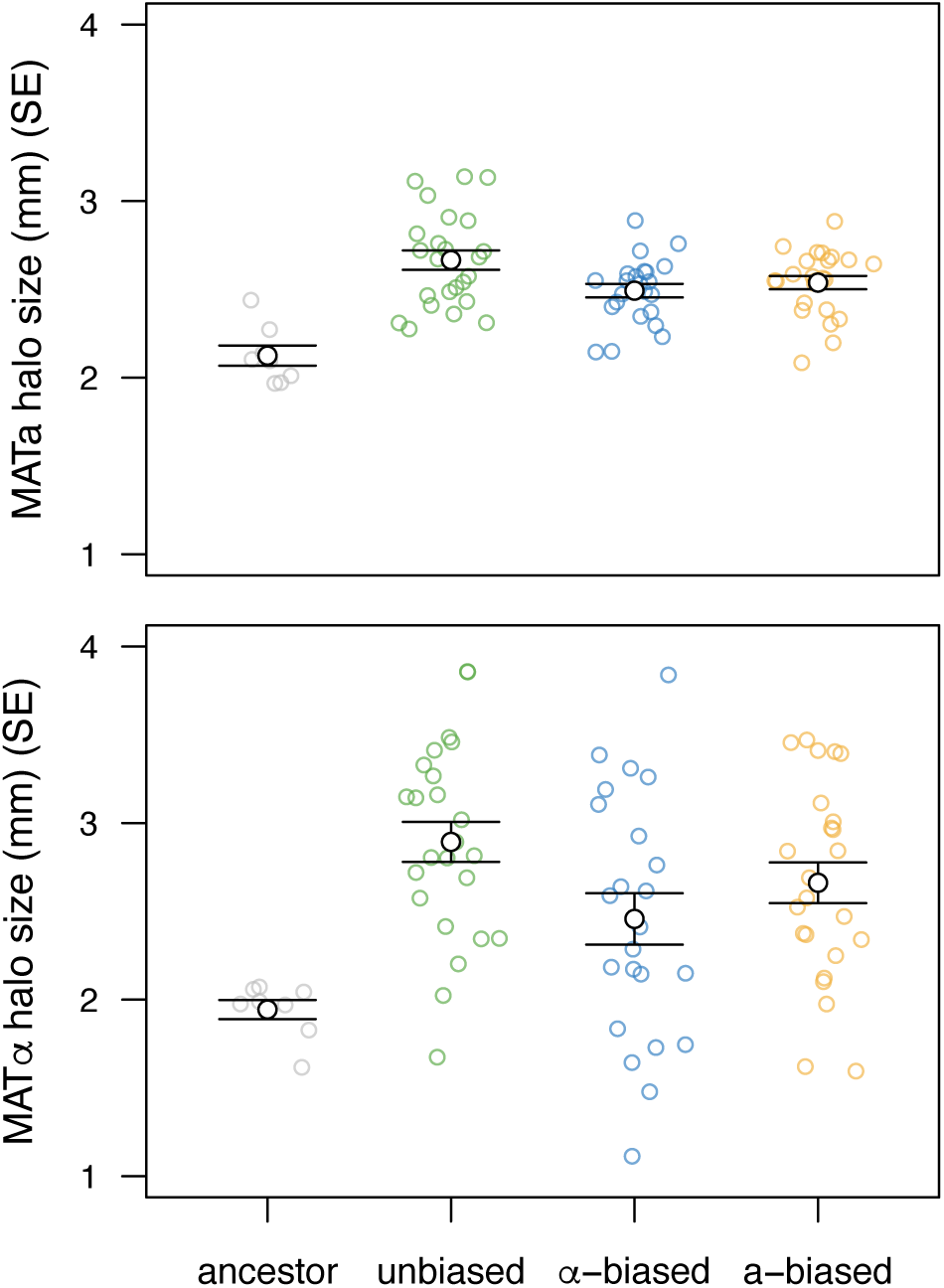
Pheromone production in evolved strains. Colored points represent mean “halo” size for each strain, with group means and standard errors shown in black. Halo size reflects the ability of a focal strain of the MATa type (top) or MAT*⍺* type (bottom) to suppress growth when plated on a lawn containing pheromone-sensitive cells of the opposite mating type.

Pheromone production rose dramatically in the evolved treatments relative to the ancestor (mixed models including evolution status: *MAT*a, *P* < 1 × 10^−8^; *MAT⍺*, *P* < 0.001). More generally, halo size differed significantly by treatment group (unbiased, a-biased, *⍺*-biased, or ancestor) for both *MAT*a strains (*P* < 1 × 10^−9^; Fig. 4) and *MAT⍺* strains (*P* < 0.001; Fig. 4). We found no evidence that halo sizes of *MAT*a and *MAT⍺* strains derived from the same experimental replicate were correlated (randomization: *P* = 0.61), except to the extent that halo sizes for both mating types increased in the evolved treatments. On average, evolved *MAT*a strains and *MAT⍺* strains induced halos that were 21.6% and 39.2% larger than their ancestors, respectively (Fig. 4), demonstrating a rapid increase in pheromone production when experimentally increasing the rate of sexual reproduction.

Accounting for differences in mean pheromone production, we found evidence that the absolute degree of mating type dimorphism was greater in the evolved populations than in the ancestor (Fig. 5A; *t* = 2.50, *P* = 0.015), though the evolved outcrossing treatments did not differ significantly from one another in this regard (*F* = 0.28, *P* = 0.76). This absolute metric reflects the magnitude of dimorphism regardless of which mating type has the higher trait value. Directional measures of dimorphism (Fig. S3A) indicated that the evolutionary increase in pheromone production was greater for MAT*⍺* cells than for MATa cells on average (see also Fig. 4), but with substantial variation among replicate populations in which mating type showed greater pheromone production.

**Figure 5.**
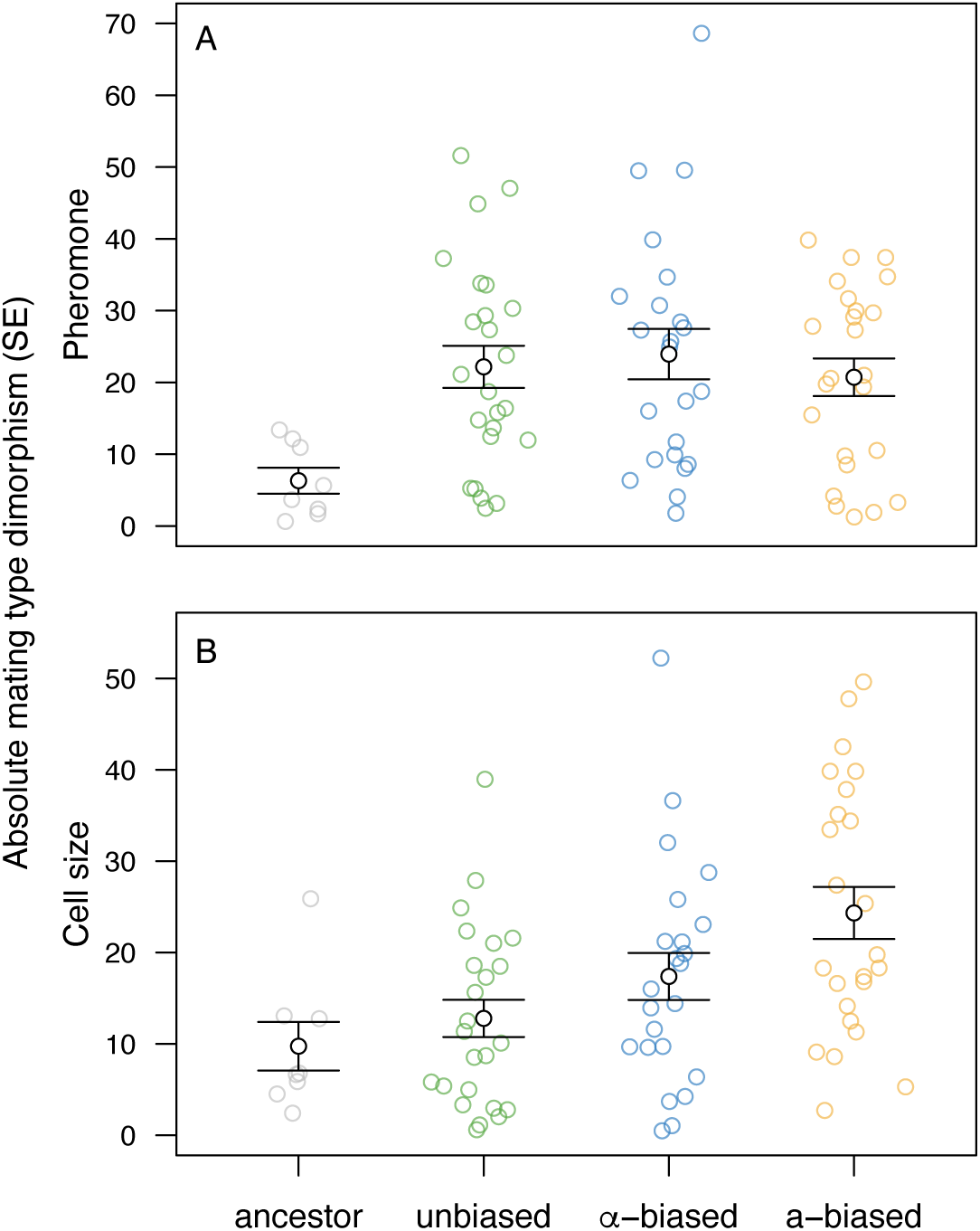
Mating type dimorphism in evolved strains. Colored points represent dimorphism between mating types of a given evolved population for pheromone production (A) and cell size (B), measured as the absolute difference between mating types, divided by the mean across mating types, expressed as a percentage. Group means and standard errors are shown in black.

### Sporulation rates

Sporulation rates (tetrad frequencies) measured from a sample of populations during the experiment were relatively stable for each mating-ratio treatment, but they declined dramatically for the selfing treatment (*t* = –7.03, *P* < 10^−5^; Fig. 6A). After the 10th sexual cycle, we retested sporulation rates for all diploid populations from the outcrossing treatments (day 3 of cycle 10), as well as the diploid ancestor (day 3 of cycle 1), using frozen cultures. These sporulation rates were all much lower (Fig. 6B) than the contemporaneous measurements (Fig. 6A), potentially due to freezing and reviving the strains. Although the sporulation rate of the unbiased mating-ratio treatment was significantly lower than the ancestor when measured at the end of the experiment (binomial GLM: *t* = –2.06, *P* = 0.04, Fig. 6B), this pattern was not seen in the contemporaneous samples (Fig. 6A). Regardless, we found no evidence for increased sporulation rates despite selection for more frequent sexual reproduction.

**Figure 6.**
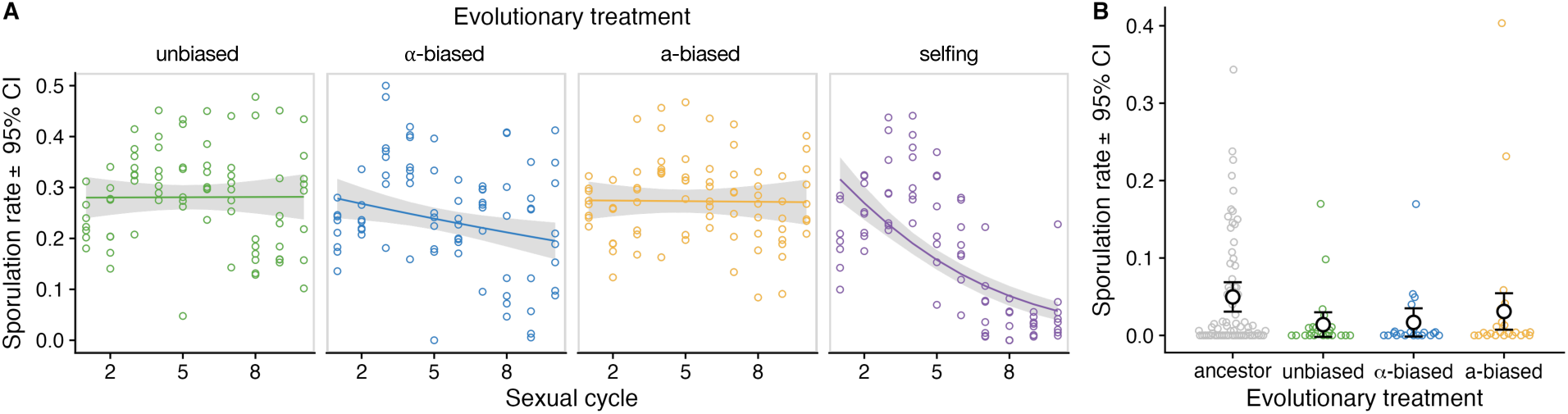
Sporulation rates. Sporulation was measured during experimental evolution (A) and at the end of the experiment (B). Each colored circle represents one hemocytometer count. Curves and grey ribbons in (A) show predicted means and 95% confidence intervals (generalized linear model). Black circles and error bars in (B) show predicted means and 95% confidence intervals (generalized linear model).

The spore staining assay confirmed that spores were absent in vegetatively growing cultures of our evolved populations (example images: Fig. 7A–C) and absent in haploid strains subjected to our sporulation protocol (Fig. 7D). We observed stained spores in diploid cultures following our sporulation protocol, generally in groups of four (Fig. 7E). In all of the selfing populations, following our sporulation protocol, we observed spores, but in groups of one or two (Fig. 7F). The apparent decline in sporulation frequency in the selfing populations over time (Fig. 6A) may therefore have been due to the production of spores in a non-tetrad formation in these populations (see below for additional results pertaining to the genotype of the selfing populations).

**Figure 7.**
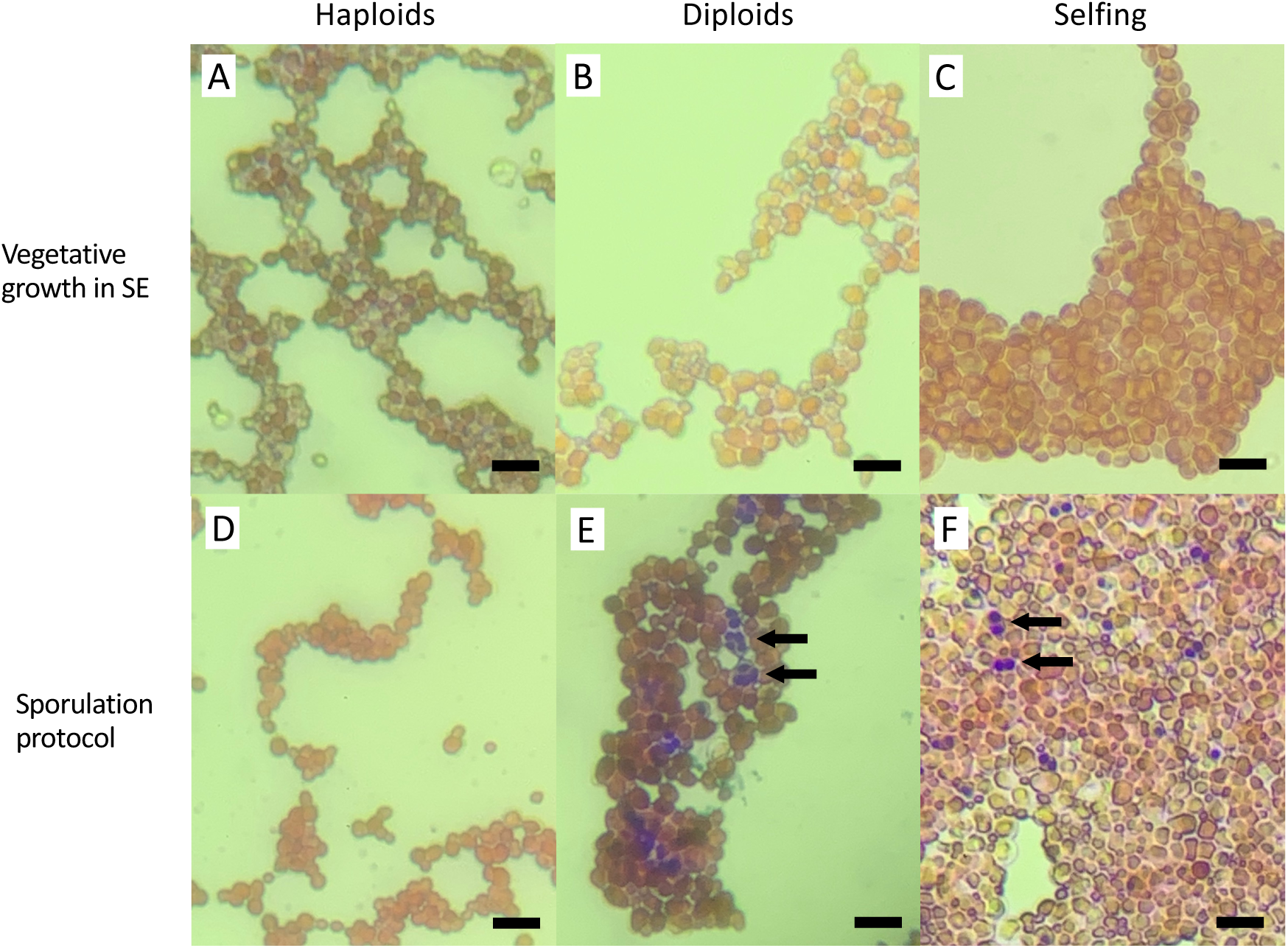
Representative spore staining images. We applied the spore straining protocol following vegetative growth in SE medium (A–C) or following the sporulation protocol (D–F) to all selfing populations (C, F), as well as haploid (A, D) and diploid controls (B, D). Spores were absent in vegetative cultures, as expected. Spores were absent in haploids following the sporulation protocol (D) and present in tetrad form in diploids following the sporulation protocol (E; arrows), as expected. In the selfing populations subjected to the sporulation protocol, spores were present singly or in groups of two (F; arrows). Scale bars are approximately 10 µm.

### Cell sizes

Using flow cytometry to measure cell sizes after the last sexual cycle (Fig. 8A), haploid cell sizes decreased relative to the ancestral haploid for both mating types and across all outcrossing treatments but did not change under the selfing treatment (*F* = 18.79, *P* < 10^−5^; contrasts: *P* < 0.001 for all comparisons with the haploid ancestor except for the selfing treatment; Fig. 8A). Across all outcrossing treatments, we found no evidence that cell sizes of *MAT*a and *MAT⍺* strains derived from the same experimental population were correlated (randomization: *P* = 0.66).

**Figure 8.**
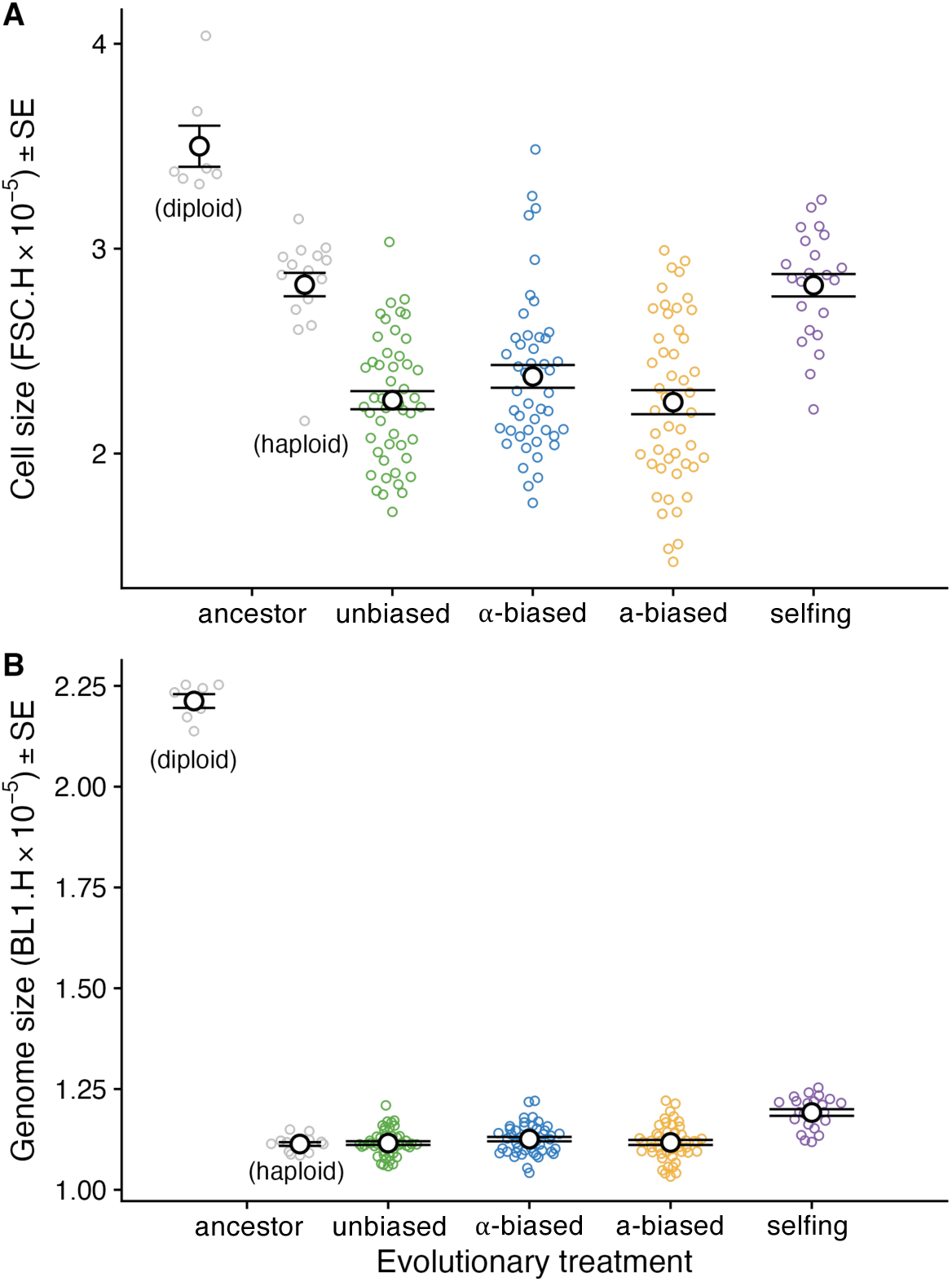
Flow cytometry data on cell size and genome size. (A) Relative to ancestral haploids, cell size in haploid cells from the outcrossing populations declined on average, but did not change in selfing populations. (B) Relative to ancestral haploids, genome size in haploid cells from the outcrossing populations remained unchanged but increased in selfing populations. Black circles and error bars indicate groupwise means and standard errors.

Accounting for differences in mean cell size, we found evidence that the absolute degree of cell size dimorphism was greater in the evolved populations than in the ancestor (Fig. 5; t = 3.02, P = 0.003). Dimorphism levels varied significantly among the outcrossing treatments (*F* = 5.46, *P* = 0.006), with greater size dimorphism present in the populations subjected to biased mating-type ratios (Fig. 5B). This absolute metric reflects the magnitude of dimorphism regardless of which mating type has the higher trait value. Directional measures of dimorphism (Fig. S3B) indicated that MATa cells were larger than MAT*⍺* cells on average, but with substantial variation among replicate populations in which mating type was larger.

### Genome sizes

Using flow cytometry to measure genome sizes after the last sexual cycle (Fig. 8B), we verified that cells grown in haploid-selective media from the outcrossing treatments retained a haploid genome content. We did not detect any difference in average genome content between the evolved and control haploids (*F* = 0.799, *P* = 0.50). While we expected the populations from the selfing treatment to be diploid, as they were sampled on day 7 following diploid selection (Fig. 1), they exhibited genome content levels similar to but slightly higher than the ancestral haploids and the outcrossing haploids (Fig. 8B; *F* = 20.19, *P* < 10^−5^). Despite showing a haploid-like genome content, our other tests (see above) indicated that cells from the selfing populations did not mate with tester strains and displayed the selectable marker phenotypes expected of diploids. Whole genome sequencing data for three selfing populations (see below) revealed approximately two-fold elevated coverage on chromosome III, where *MAT* is located (Fig. S4). Using PCR, we verified the presence of both MATa- and MAT*⍺*-specific sequences in all of the selfing populations (Fig. S5). Together, these observations indicate that the selfing populations became haploid during the experiment but retained two copies of some or all of chromosome III (disomy), with a MATa/MAT*⍺* genotype that included each drug resistance marker; these aneuploid selfing lines remained capable of spore production and survived zymolyase treatment, despite not producing tetrads (Fig. 7).

### Trait correlations

We found a significant negative correlation between cell size and growth rate in the selective medium in the outcrossing treatments (mean *r_S_* = –0.182, randomization *P* = 0.023; Fig. S6). This pattern was driven by the biased mating ratio treatments, which showed a more negative correlation than the unbiased treatment (randomization *P* = 0.007; Fig. S6). Specifically, faster growth was associated with smaller cell size among the populations that evolved under biased mating ratios.

Cell size in haploids could affect the performance of diploids resulting from mating (Smith et al. 2014). We found that the growth rate of diploids in permissive media (the condition of mating and initial diploid growth) was positively correlated with the average cell size of the constituent haploids (mean *r_S_* = 0.258, randomization *P* = 0.031; Fig. S7). The size and mating rate of haploids was not significantly correlated with sporulation frequency in the resulting diploids; however, we found suggestive evidence that strains with faster growth in diploid-selective media also had reduced sporulation frequency (mean *r_S_* = –0.219, randomization *P* = 0.064).

Averaging across the outcrossing treatments, we found that cell size was positively correlated with mating rate under the evolved mating-type ratio assay conditions (mean *r_S_* = 0.203, randomization *P* = 0.013; Fig. S8). Because of this relationship between cell size and mating rate, we hypothesized that strains reproducing faster––but generating smaller cells––might experience a trade-off with reduced mating competitiveness, especially for the common mating type experiencing stronger sexual selection. In the lines evolved under biased mating-type ratios, we found a significant negative correlation between growth rate of the common cell type in selective media and the average mating rate across assay conditions (mean *r_S_* = –0.39, randomization *P* = 0.013; Fig. S9), whereas there was no significant correlation for the rare cell type (mean *r_S_* = 0.10, randomization *P* = 0.49; Fig. S9). This difference in correlation patterns between the common versus rare type is unlikely to occur by chance (randomization *P* = 0.016). In populations from the unbiased outcrossing treatment, we did not detect any correlation between growth and mating (mean *r_S_* = –0.10, randomization *P* = 0.66; Fig. S9).

### Genomic variation

We sequenced a subset of the experimental populations to gain insight into the molecular basis of phenotypic change. As noted above, our examination of sequencing coverage revealed two-fold excess coverage on chromosome III in the three selfing populations sequenced; combined with our genome size data, this indicates the selfing populations were haploid but disomic for chromosome III. In the samples from outcrossing populations, we identified 81 non-synonymous variants affecting 64 different genes. 31 variants were found in more than one sample. Three of the affected genes are known to be involved in the mating process. First, in one of the MAT*⍺* samples from the *⍺*-biased treatment, we identified a missense mutation of moderate impact in *GPA1* (YHR005C): E51Q. This gene encodes the *alpha* subunit of the G protein in the G-protein-coupled pheromone receptor; null mutants have impaired mating response (Chasse et al. 2006) and decreased sporulation efficiency (Deutschbauer et al. 2002); reduction of function mutants lack pheromone-induced cell cycle arrest (Fujimura 1989). The E51Q mutation may have thus been a gain-of-function mutation leading to an increased mating rate for this line.

Second, we identified a missense mutation of moderate impact in *SAG1* (YJR004C): F336V. This variant was shared among 11 of the 14 sequenced samples, including two of the three selfing strains (the third inconclusive due to low coverage), and absent in the ancestor. This gene produces the alpha-agglutinin of *MAT*α-cells; null mutants fail to agglutinate in response to MATa-factor (Doi et al. 1989). This variant could influence mating efficiency by altering the properties of cell adherence during conjugation.

Third, we found two variants in *CCW12* (YLR110C), a mannoprotein that localizes to newly synthesized cell walls, including the mating projection tip (“shmoo”). Null mutants of this gene have abnormal shmoo morphology, increased mating-related mortality (Ragni et al. 2011), and decreased agglutination and mating efficiency (Mrsa et al. 1999). One of the variants was an A123S missense mutation of moderate impact, which we found in three separate evolutionary replicates (two MATa-biased populations and one MAT*⍺*-biased population). The second variant was a Y110C missense mutation of moderate impact in one of the MAT*⍺*-samples from the a-biased treatment. These variants could influence mating efficiency by altering the cell wall properties of mating projections.

We also found variants in three different genes involved in the sporulation process. *RDH54* (YBR073W) is involved in recombinational repair and supercoils DNA. Null and reduction-of-function mutants of this gene have decreased sporulation rate and spore viability (Chi et al. 2006). We found a V668E missense mutation of moderate impact in this gene in a MATa sample from the *⍺*-biased treatment, affecting the predicted helicase domain. We also found an E82G missense variant of moderate impact in *ATG9* (YDL149W) shared by the MATa and MAT*⍺* samples of an evolutionary replicate. *ATG9* is a lipid scramblase involved in autophagy. Both null mutants and reduction of function mutants of this gene are unable to sporulate (Tsukada and Ohsumi 1993; Lang et al. 2000; Enyenihi and Saunders 2003). Additionally, we found an in-frame deletion of 36 bp (amino acids 138-150) in the middle of *TIR1* (YER011W), which encodes a cell wall mannoprotein. This variant was found in six of our sequenced samples, two of which are MATa and MAT*⍺* samples from the same evolutionary replicate. Null mutants of *TIR1* have decreased sporulation efficiency (Marston et al. 2004). While complete deletion of each of these genes leads to reduced sporulation, we speculate that the variants we observed could have subtler effects, serving to improve or maintain sporulation performance and spore viability under the conditions of our experiment.

As described further in S7 Text, we identified other variants of potential interest, several of which were associated with the auxotrophic selection used in our experiment.

We additionally examined coverage profiles for each chromosome in the sequenced populations to identify segmental duplications or deletions. We discovered a duplication of approximately 153 kb on chromosome VIII in a MAT*⍺* sample from an outcrossing *⍺*-biased population. The duplicated region is in the middle of the right arm of the chromosome, with approximate breakpoints at positions 228965 and 382489 (Fig. S10). This region includes 79 genes, of which at least six are potentially relevant to sexual traits: *SAE3* (involved in meiotic recombination), *STE12* (transcription factor activating genes involved in mating response to pheromone), *NAM8* (involved in meiosis and spore formation), *KIC1* (localizes to the mating projection), *NDT80* (transcription factor required for meiosis and sporulation), and SPS100 (required for spore wall maturation).

## Discussion

In this study, we maintained yeast populations with frequent rounds of sex under different mating regimes (Fig. 1). The mating treatments included conditions promoting outcrossing with either equal proportions of the two mating types or a skewed ratio in favor of one mating type (*⍺*-biased or a-biased), and a treatment promoting within-tetrad mating soon after germination (selfing). Our protocol forced cells to reproduce sexually once in every 64 rounds of cell division, which we predicted would impose stronger selection for sexual characteristics than experienced by wild yeast (Ruderfer et al. 2006; Magwene et al. 2011).

### Evolution of mating characteristics

We found an increase in mating rate in all treatments compared to the ancestor when assayed with an equal ratio of the two mating types, confirming that the response to selection for mating was strong in our experiment, with no significant differences between evolutionary treatment groups (Fig. 3A). When tested under *⍺*-biased conditions, on the other hand, the lines that evolved under that condition mated at a higher rate than the other two treatment groups, a sign of adaptation to the operational mating-type ratio (Fig. 3B). Furthermore, we found that it was the common mating type (in this case *MAT⍺*) that evolved to be better at mating (Fig. S2), consistent with this mating type experiencing stronger competition. In contrast, the mating rate in the a-biased ratio was low in all treatment groups (Fig. 3C), and we did not find evidence that populations from the a-biased treatment performed best in this scenario (Fig. S2). Similarly, while both mating types increased pheromone production under experimental evolution, the increase was greater in *MAT⍺* cells (Fig. 4).

A potential factor accounting for the difference in evolutionary responses of the MATa and MAT*⍺* cells is the Bar1p protease produced by *MAT*a cells (Sprague and Herskowitz 1981; Anders et al. 2021). This extracellular enzyme digests *⍺* factor (the mating pheromone produced by *MAT⍺* cells). Bar1p ensures that the *MAT*a cell does not initiate mating unless it is in close proximity to a *MAT⍺* cell, *i.e.*, when the concentration of *⍺* factor is high. When *⍺* factor reaches and remains in the pheromone receptors of a *MAT*a cell, the cell stops dividing and prepares for mating. In liquid culture, the ratio of *MAT*a and *MAT⍺* cells affects the relative concentrations of Bar1p and *⍺* factor and, hence, the propensity of *MAT*a cells to mate (Banderas et al. 2016; Anders et al. 2021). The abundance of Bar1p in the a-biased mating ratio may have shifted the balance, digesting the rare *⍺* factor in the culture, preventing rapid mating (Fig. 3C). By allowing *MAT*a cells to continue vegetative growth, Bar1p may have skewed the sex ratio in a-biased treatments back to 50:50. A mutation in *MAT*a cells that decreased production of Bar1p would not increase its mating rate, as the effect of a single cell on the amount of enzyme in the medium would be negligible. Conversely, *MAT⍺* cells in the a-biased treatment that increased pheromone production might have improved chances of mating, explaining the larger halo sizes observed (Fig. 4).

In contrast to the extracellular Bar1p protease produced by *MAT*a cells, the corresponding enzyme in *MAT⍺* cells that degrades a-factor, Afb1p, remains cell-associated [see (Steden et al. 1989; Marcus et al. 1991) for early studies and (Huberman and Murray 2013) for isolation, characterization, and mapping of the Afb1 protein]. Consequently, mutations in this or other genes that increase sensitivity of *MAT⍺* to a-factor could allow earlier mating than competing *MAT⍺* cells. The difference in localization between Bar1p and Afb1p could thus underlie our observation that *MAT⍺* cells, but not *MAT*a cells, evolved to be better mating competitors when common.

By whole-genome sequencing of a subset of lines, we obtained evidence for several variants with potential effects on the expression of mating pheromone (to overcome the degradation of Bar1p and Afb1p) and mating pheromone receptors (to increase sensitivity to the presence of the other mating type). In addition to selection for more efficient sexual reproduction, the auxotrophic and antibiotic markers used to maintain the alternation of haploid and diploid phases (McDonald et al. 2016) would have also been under strong selection during our experiment. Because *LEU2* is under the same promotor as the *STE3* receptor used by *MAT⍺*, a change in their transcription factors could have favored both a higher growth rate in leucine-limited medium and an increased mating rate. This cannot be the only explanation for the increase in mating rate, however, because we do not see similar increases in mating rate in the other outcrossing treatments groups (that had similar selection for *MAT⍺* growth with *LEU2*). Furthermore, it fails to explain the negative correlation we found between mating rate and growth rate in the common mating type in the biased mating ratios (Fig. S9).

### Evolution in other traits

Growth rates increased under experimental evolution (Fig. 2), which is likely due at least in part to adaptation to the lab conditions and haploid-selective media, rather than adaptation to repeated rounds of sexual reproduction. Although we saw dramatic increases in growth rates (Fig. 2) and mating rates (Fig. 3), we observed little change in sporulation rates in the outcrossing treatments (Fig. 6). Because *S. cerevisiae* is known to mate readily but show low sporulation propensity (especially in lab strains), we had expected strong selection for increased sporulation as a “weak link” in the sexual cycle. Studies on both natural isolates (Gerke et al. 2006) and laboratory strains (Codón et al. 1995; Deutschbauer et al. 2002; Enyenihi and Saunders 2003; Deutschbauer and Davis 2005; Ben-Ari et al. 2006; Hall and Joseph 2010) suggest that many loci are involved in determining sporulation efficiency and that mutations arise that both increase and decrease sporulation rate. In addition, increased sporulation rates in response to frequent rounds of sex have been shown to evolve in other experimental studies with yeast (Zeyl et al. 2005; Thomasson et al. 2021). Interestingly, another study that evolved populations under mating competition also saw decreases in sporulation rate (Reding et al. 2013), suggesting a potential trade-off between traits that we selected for simultaneously.

Correlations between traits in the evolved populations provided additional insight into the possible trade-offs involved in adaptation to conditions of increased sexual competition. First, we observed a trade-off between growth rate (in the haploid-selective medium) and cell size in haploids, particularly in the biased mating-ratio treatments (Fig. S6). Second, we saw that higher mating rates were associated with lower growth rates, but only for the common mating type in the biased treatments (Fig. S9). These results suggest that two alternative evolutionary strategies may have been available for the mating type experiencing stronger sexual selection: either gain a numerical advantage over competitors through faster growth at the expense of cell size and mating rate (i.e., those lines with higher growth rates but lower per-capita mating rates and smaller cells) or increase attractiveness to potential mating partners (i.e., those lines with lower growth rates but high per-capita mating rate and larger cells). While a trade-off between mating rate and growth rate has been reported previously (Lang et al. 2009), our study suggests increasing either trait may be an appropriate response to increased sexual selection, at least initially. Of these strategies, an increase in gamete numbers at the expense of gamete size appears to result in reduced growth of the diploids that are formed by those gametes (Fig. S7), which is a key trade-off in models of the evolution of anisogamy (Parker et al. 1972; Randerson and Hurst 2001; Parker 2011).

The evolution of cell size in one mating type could encourage compensatory evolution in the opposite mating type. Our outcrossing populations evolved increased mating type dimorphism for cell size and pheromone production (Fig. 5), suggesting that sexual competition imposes divergent selection pressures on the two mating types and that the mating types can evolve independently to some degree. However, this dimorphism was not strongly directional across replicates in any given treatment group (Fig. S3), and we did not observe significant correlations in cell size between mating types, suggesting that antagonistic coevolution between mating types may require more evolutionary time than mating-type specific evolution. Further experimental evolution of yeast under strong sexual selection may therefore shed light on the evolution of gamete-size dimorphism (anisogamy).

### MATa/*⍺* haploids evolved repeatedly under selection for selfing

In the selfing treatment that allowed for mating immediately after germination, all of the replicate populations evolved a genome size typical of haploids (Fig. 8B) while retaining both mating types. Our sequence data for three selfing populations revealed disomy of chromosome III in all cases; this may be the most accessible route for haploid cells to retain both *MAT* isoforms, providing an advantage by avoiding mating altogether. We also found that these strains were capable of sporulation, producing spores in pairs rather than tetrads. *MAT*a/*⍺* haploids with disomy-III have been shown to successfully sporulate, producing two spores (dyads), only if they also carry mutations that allow meiosis I bypass; the resulting spore cells are typically *MAT*a/*⍺*, but sometimes lose a copy of chromosome III (Wagstaff et al. 1982). In experimental evolution studies with *S. cerevisiae*, which is typically diploid in nature (Peter et al. 2018; Loegler et al. 2024), it is more common to observe transitions from haploidy to diploidy than the reverse (Gerstein et al. 2006; Gerstein and Berman 2015; Fisher et al. 2018; Harari et al. 2018; Gerstein and Sharp 2021), although transitions to haploidy have been observed [e.g., see (Crandall et al. 2023) for a recent example]. In our experiment, sporulating *MAT*a/*⍺* haploids were able to survive clonally through the entire selfing selection regime: they created spores resistant to zymolyase treatment, and they retained both drug resistance markers, allowing survival in the part of our protocol meant to select for diploids (Fig. 1A). Additionally, *MAT*a/*⍺* haploids avoid costly haploid-specific expression (Lang et al. 2009; Haber 2012), as well as the need for mating, which could provide a growth advantage compared to the ancestor during germination. Finally, this case illustrates that meiosis (or a meiosis-like process) and sporulation can be uncoupled from ploidy cycles.

### Evidence for additional genomic changes

Although we sequenced only a subset of the experimental populations, we found evidence for molecular evolution in multiple genes related to growth, mating and sporulation, including apparent cases of convergence. We also found a large duplication that increased the copy number of multiple genes with roles in sexual reproduction. While confirming the role of any particular variant would require further work to compare alternative alleles on a standard genetic background, in aggregate, these data indicate that there is ample molecular genetic variation to allow for rapid and multifaceted responses to selection for sexual performance in yeast.

### Conclusion

We forced yeast to go through alternating rounds of asexual and sexual reproduction and manipulated the opportunity for sexual selection. We detected evolutionary responses in growth, cell size, pheromone production, and mating. Despite strong selection for meiotic capacity, the propensity to sporulate showed little evolutionary change in our experiment. The trait correlations we observed suggest that the common mating type can either evolve to grow more rapidly, providing a numerical advantage during mate competition at the expense of cell size and diploid fitness, or can evolve to become more attractive to the opposite mating type, despite a slower growth. This finding supports the idea that disruptive selection on gamete traits can promote the evolution of anisogamy (Parker et al. 1972; Parker 2011).

We found that *MAT⍺* cells responded more strongly under experimental evolution, evolving to become more efficient at mating, and we hypothesize that this asymmetry may be caused by biological differences in the way *MAT*a and *MAT⍺* cells respond to the presence of pheromones of the opposite mating type. Such asymmetry in gamete signaling may be a necessary feature of single-celled eukaryotic mating systems (Hadjivasiliou and Pomiankowski 2016). Thus, while *S. cerevisiae* is isogamous at a morphological level, this species displays mating type dimorphism at molecular and functional levels, affecting evolutionary responses to sexual selection.

## Data availability

Raw data and scripts used for analyzes are available at doi:10.6084/m9.figshare.30266410. Whole genome sequencing data will be available on Sequence Read Archive (www.ncbi.nlm.nih.gov/bioproject/1337161) pending publication.

## Acknowledgements

We are grateful for laboratory assistance from Stephan Koenig, Mathew Stasiuk, Christina Hsu, Kismet Somal, Bryn Wiley, and Penelope Kahn. We also thank Audrey Gasch for providing pheromone sensitive strains and Auguste Dutcher for helpful discussion. This research was supported by the Natural Sciences and Engineering Research Council of Canada Discovery Grant to SPO (RGPIN-2022-03726) and by the National Institute of General Medical Sciences of the National Institutes of Health award number R35GM154954 to NPS.

## Supplementary Material

### S1 Text. Strain details and selective markers

Because the STE5 promoter is active only in haploids, only haploids can grow on uracil drop-out medium. Haploids can also be selected against by the use of 5-fluoroorotic acid (5-FOA), which is converted into a toxic byproduct by the uracil biosynthesis pathway. This means only diploid yeast are able to grow on medium supplemented with 5-FOA. Because the STE2 promoter is active only in MATa haploids, growth on histidine drop-out medium selects for MATa haploids. Similarly, the STE3 promoter is MATα-specific, giving these cells the ability to grow on leucine drop-out medium. The antibiotic markers KanMX and HphMX have been inserted close to the MAT locus. These markers select for MATa through resistance to geneticin (g418) and for MAT⍺ through resistance to hygromycin, respectively. When the two antibiotics are combined in the medium, only MATa/MAT⍺ yeast can grow. Spot assays of the haploid lines used in this study, as well as the diploid made from mating the two, on the three different selective media used in this study is shown in Fig. S1. Because the *MAT⍺* strain received had acquired a petite phenotype, we crossed the *MAT*a and *MAT⍺* strains, sporulated the resulting diploid, and obtained haploids with normal respiratory function through tetrad dissection. This has the added advantage of re-mixing the genetic backgrounds of the two mating types.

### S2 Text. Dilutions for mating

For all dilutions, we used dilution factor, *DF = V_f_ / V_i_*, where *V_i_* is the volume of culture and *V_f_*, is the total volume of medium with added culture. In the equal mating ratio treatment, we added 100 μL of saturated culture of each mating type to the well (*V_i_ =* 100), added and removed 800 μL of medium. The 100 μL of the opposite mating type culture contributes to the total volume and gives *V_f_* = 1000 (DF 10). Then, we added 800 μL of medium (*V_i_ =* 200, *V_f_ =* 1000, DF 5). This means, for each mating type, there was a 50-fold dilution (10×5), while for cell concentration, there was a 25-fold dilution (5×5). In contrast, in the skewed mating ratio we used 1 mL of culture of the one mating type and 100 μL for the other, then removed 900 μL (a 11-fold dilution for the rare mating type, *V_i_ =* 100, *V_f_ =* 1100). We then added 800 μL of medium (*V_i_ =* 200, *V_f_ =* 1000, DF 5). This resulted in a 55-fold dilution of the rare mating type and a 5.5-fold dilution for the common mating type (roughly as in the equal mating ratio treatment). There was a 5-fold dilution for overall cell concentration.

### S3 Text. Mating assays with partners from different populations

We chose *MAT*a and *MAT⍺* haploid isolates from the three populations exhibiting higher than average mating rates when tested in the mating ratio under which they had evolved (a-biased or *⍺*-biased). We combined the six *MAT*a populations and six *MAT⍺* populations in a total of 36 combinations, measured mating rate three times, split over two blocks, under both biased mating ratios (a-biased or *⍺*-biased). Because we had observed a higher mating rate in the *⍺*-biased ratio in our previous assays, we diluted these mating cultures 10-fold more than the cultures mated in the a-biased ratio. Because the mating rate was very low, we decided to exclude measurements from plates where the maximum number of diploids that could have been plated on the selective plates (D_max_/the dilution factor used for the selective plate) was below 50. To test whether the increase in mating rate of the couples was achieved by adaptation to high competition of the more common mating type (*MAT*a cells in the a-biased treatment and *MAT⍺* cells in the *⍺*-biased treatment), we fit a linear mixed-effect model with mating couple as a random effect and with assay mating ratio, the treatment under which the *MAT*a population evolved, and the treatment under which the *MAT⍺* population evolved as fixed effects. We allowed for three-way interactions in this model. We assess the effect of the interactions with pairwise comparisons (conditional on one of the three factors) using the *emmeans* package. One of the MAT*⍺* populations we had chosen proved to be contaminated with a selfing population and so was excluded from the analysis.

### S4 Text. Spore staining protocol

1. Transfer 30 μL of yeast culture to a glass slide.
2. Heat the slide gently to allow liquid to evaporate.
3. Fix cells by passing the slide through a Bunsen burner flame for 1 s, repeated three times.
4. Return slide to a warm hotplate and add 1-2 drops of aniline crystal violet [Solution 1].
5. Warm slide for 3 min, then gently rinse with distilled water.
6. Add 2-3 drops of de-staining solution [Solution 2] to remove excess dye.
7. Gently rinse with distilled water and inspect under magnification; if deep purple coloration remains, repeat steps 6 and 7.
8. Shake gently to remove excess water and add 1-2 drops of safranin O [Solution 3].
9. Wait at least 15 s, then remove liquid with a wipe or allow it to evaporate.
10. Examine slide under magnification; vegetative cells appear red, and spores appear blue or purple.

<u>Solution 1</u>: 5 g crystal violet powder (Aldon Corp SE; CAS no. 548-62-9; VWR cat. no. 470300-948); 10 mL 95% ethanol (general lab-grade); 2 mL aniline 99.5% (Sigma-Aldrich; CAS no. 62-53-3; cat. no. 242284-100ML); 20 mL distilled water.

<u>Solution 2:</u> 6 mL HCl (general lab-grade); 190 mL 100% ethanol (general lab-grade); 10 mL distilled deionized water.

<u>Solution 3</u>: Safranin O, 0.25% aqueous (Aldon Corp SE; CAS no. 477-73-6; VWR cat. no. 470302-316).

### S5 Text. Flow cytometry

For each strain, we inoculated 1 mL of YPAD medium with 10 μL of freezer stock in 96-well plates and incubated with shaking overnight. We transferred 20 μL of the overnight cultures 1.5 mL microcentrifuge tubes and washed in 1 mL of sterile water before killing cells using 1 mL of cold 70% ethanol. We kept the tubes either at room temperature for at least 1 h, or overnight in the refrigerator. After the ethanol treatment, we pelleted cells at 2500 rpm for 2 min, then washed in 1 mL of 50 mM sodium citrate, then treated with RNAase (25 μL of 10 mg/mL RNAase added to 1 mL of sodium citrate). We kept the samples in the RNAase mixture overnight at 37 C. The following day, we pelleted cells, removed supernatant, and stained with Sytox Green (30 μL of 50 mg/mL Sytox Green added to 1 mL of sodium citrate). We left the stained cultures at room temperature and in darkness overnight.

### S6 Text. Phenotyping

Following experimental evolution, we spotted cultures on SE and selective plates: SE + 5FoA + g418 + hygromycin for diploid, SE –Ura –His + g418 for MATa haploids and SE –Ura – Leu + hygromycin for MAT⍺ haploids. After two days of growth at 30 C we scored the plates visually. Two lines showed growth on more than one kind of selective medium: one MAT⍺ haploid (strain 45) showed weak growth on the diploid selective medium, and one line from the selfing treatment (strain 177) showed growth both on the diploid selective and on the ⍺-selective medium. We did not further investigate whether these rare exceptions in growth were due to polymorphism within the cultures, mutational effects, or experimental error.

To test mating type, we first spotted strains on YPAD, and replica plated onto new YPAD plates for mating to the MAT⍺ and MATa tester (both *his1-123*), which were grown as lawns. After 6 h, we replica plated mating plates onto media lacking eight amino acids (SC –Arg – His –Leu –Lys –Met –Trp –Ade –Ura) and incubated for 2 d. While the plates that resulted from mating to the MATa tester were easy to score and grew according to expectations, the plates resulting from mating to the MAT⍺ tester showed contradictory results (both in comparison to the results from the growth on selective plates as well as mating to the MATa tester), suggesting contamination of the tester stock. Because of uncertainty about the MAT⍺ tester stock used, we only report results from the MATa tester stock.

### S7 Text. Additional genetic variants

We identified genomic variants in genes related to the auxotrophic markers we used for selection. One MATa sample from the a-biased treatment had a complex variant in *LEU2*, constituting an in-frame addition of 38 amino acids following protein position 17. In our experiment, *LEU2* was under control of the *STE3* promotor, which codes for the a-factor pheromone receptor, expressed only in MAT*⍺* cells. We also found two insertions in *HIS3* causing frameshifts, starting at amino acid positions 14 and 16, one affecting MATa and MAT*⍺* samples from the same population from the *⍺*-biased ratio treatment, and another affecting a MATa sample from the a-biased treatment. In our experiment, HIS3 was under control of the STE2 promotor, which codes for the *⍺*-factor pheromone receptor, expressed only in MATa cells. Three different populations also had missense mutations of moderate impact in URA2 (whereas URA3 was under the control of a haploid-specific promotor in our experiment), with both the MATa and MAT*⍺* samples affected in two of those cases, and just the MAT*⍺* sample affected in the remaining case. In the selfing populations we sequenced, we identified three variants in genes implicated in growth: T229I in SIS2 (YKR072C), Y329N in MYO5 (YMR109W) and Y391* in CAF120 (YNL278W). In those cases in which a mutation would putatively disrupt function in a gene essential for growth in our experiment, we speculate that these mutations could have occurred during the period of growth in non-selective media immediately prior to DNA extraction.

## Supplementary Figures and Tables

**Figure S1.**
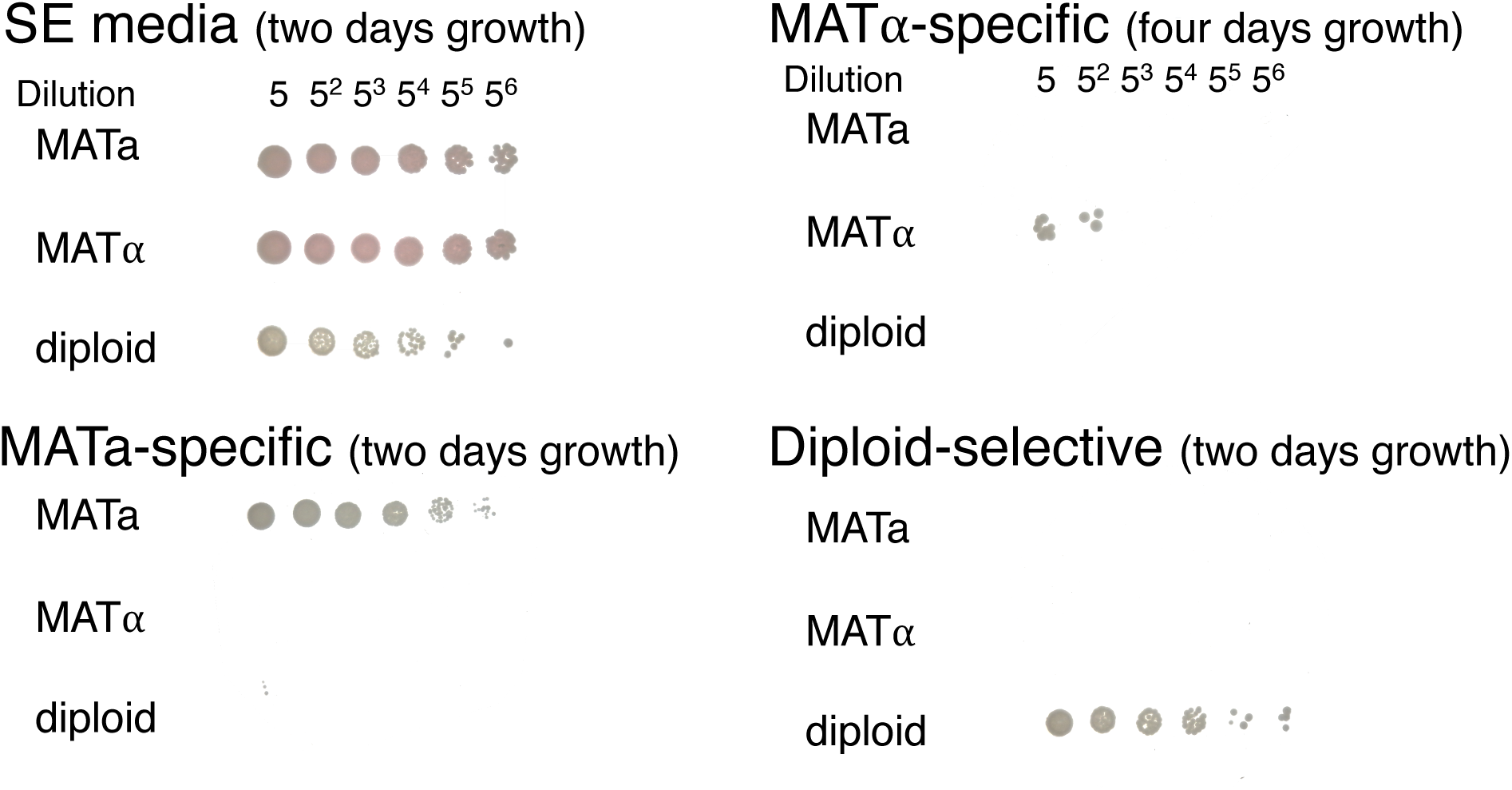
Spot assay of ancestral lines on selective media. Spots represent the growth of yeast from each ancestral strain type on solid media.

**Figure S2.**
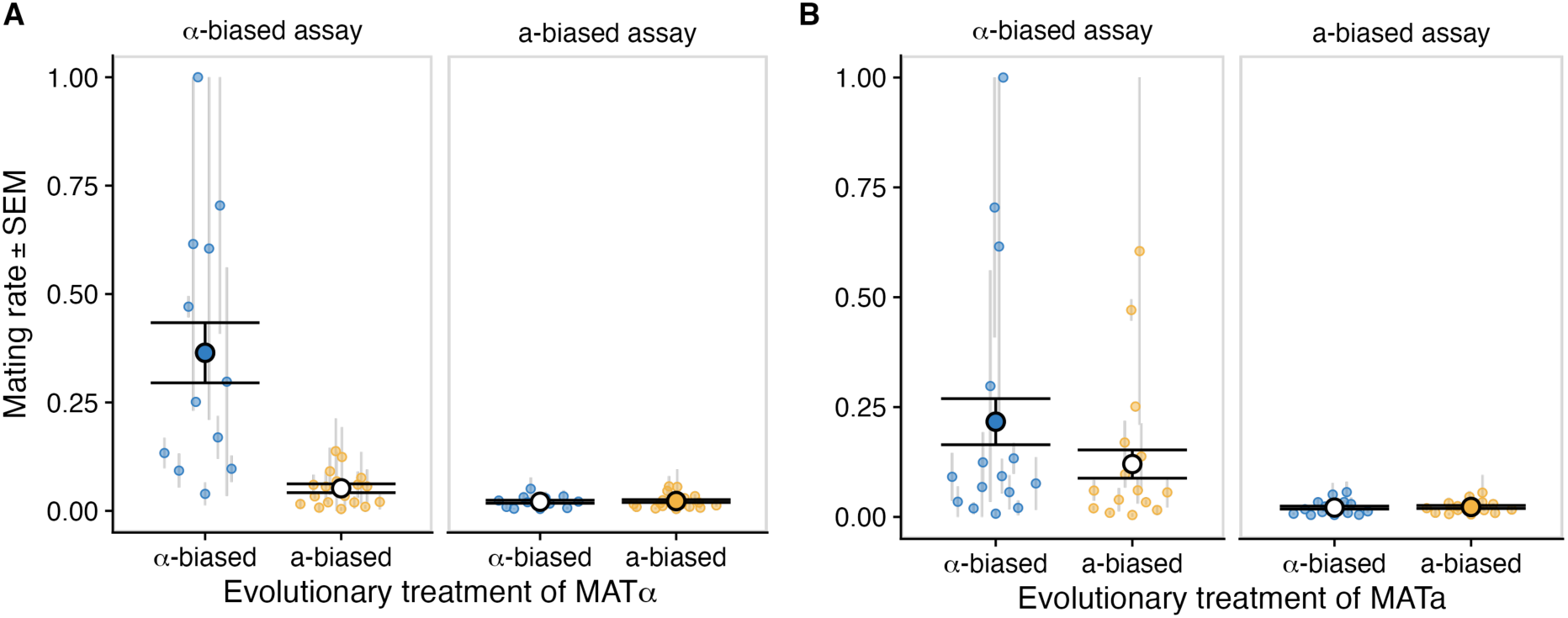
Cross-couple mating rates under alternative mating-type ratios. Panels indicate the assay ratio and the evolutionary treatment of the *MAT*⍺ line (A) or *MAT*a line (B) used in the cross. The mating rate of a cross was measured at least twice. Points represent means and grey lines represent standard errors. Black circles and error bars show group means and standard errors, with the circle filled for the treatment group that evolved under the assayed condition.

**Figure S3.**
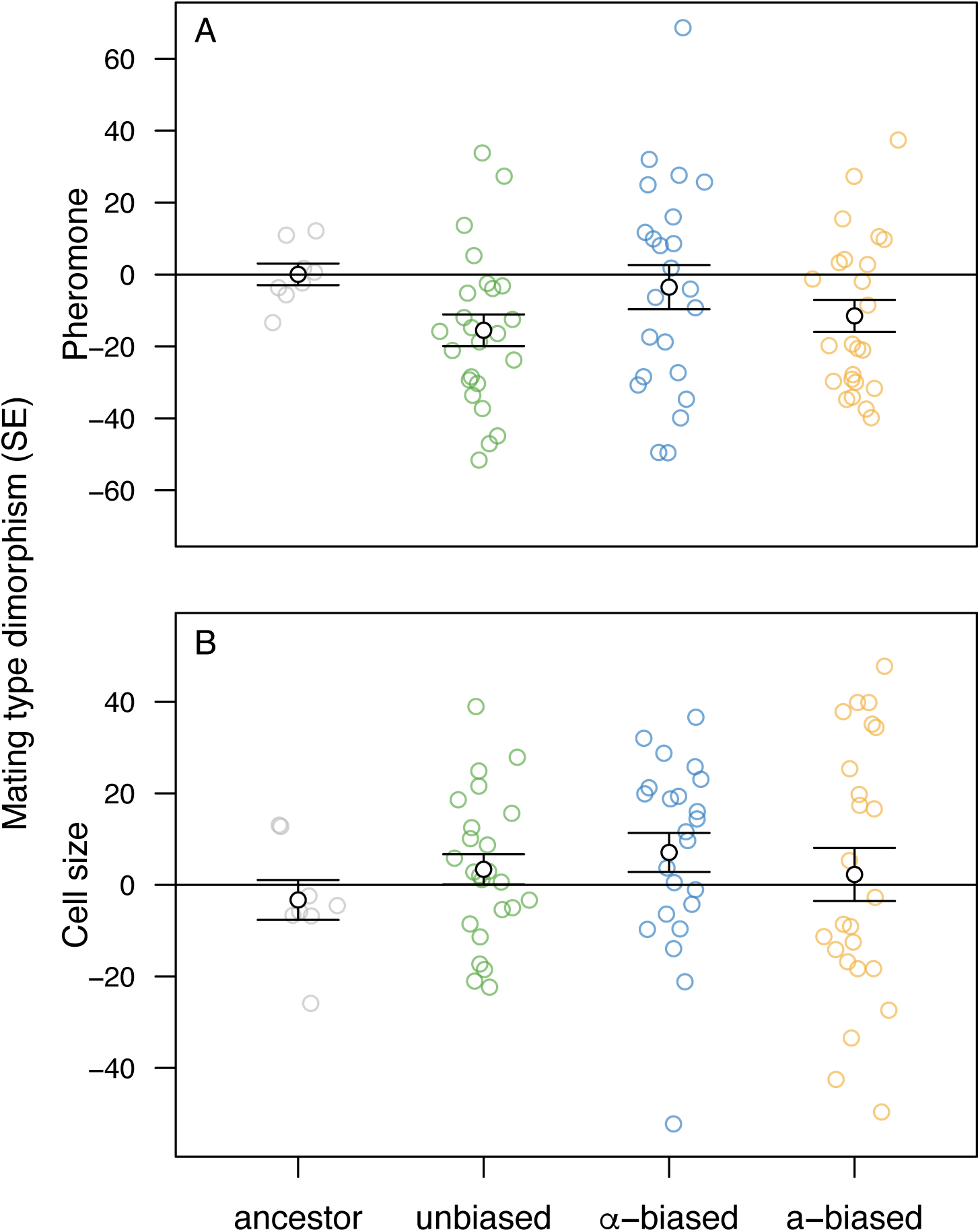
Directional measures of mating type dimorphism. Each point represents the difference in trait values between mating types for a given replicate population (MATa – MAT*⍺*). Open black circles represent group means. (A) Following evolution, pheromone production increased in both mating types (see Fig. 5), but the change was greater for MAT*⍺* on average. (B) Following evolution, haploid cell size decreased on average (see Fig. 9A), but MATa cells tended to be larger than MAT*⍺* cells. For both traits, the absolute level of mating type dimorphism increased under experimental evolution (see Fig. 6).

**Figure S4.**
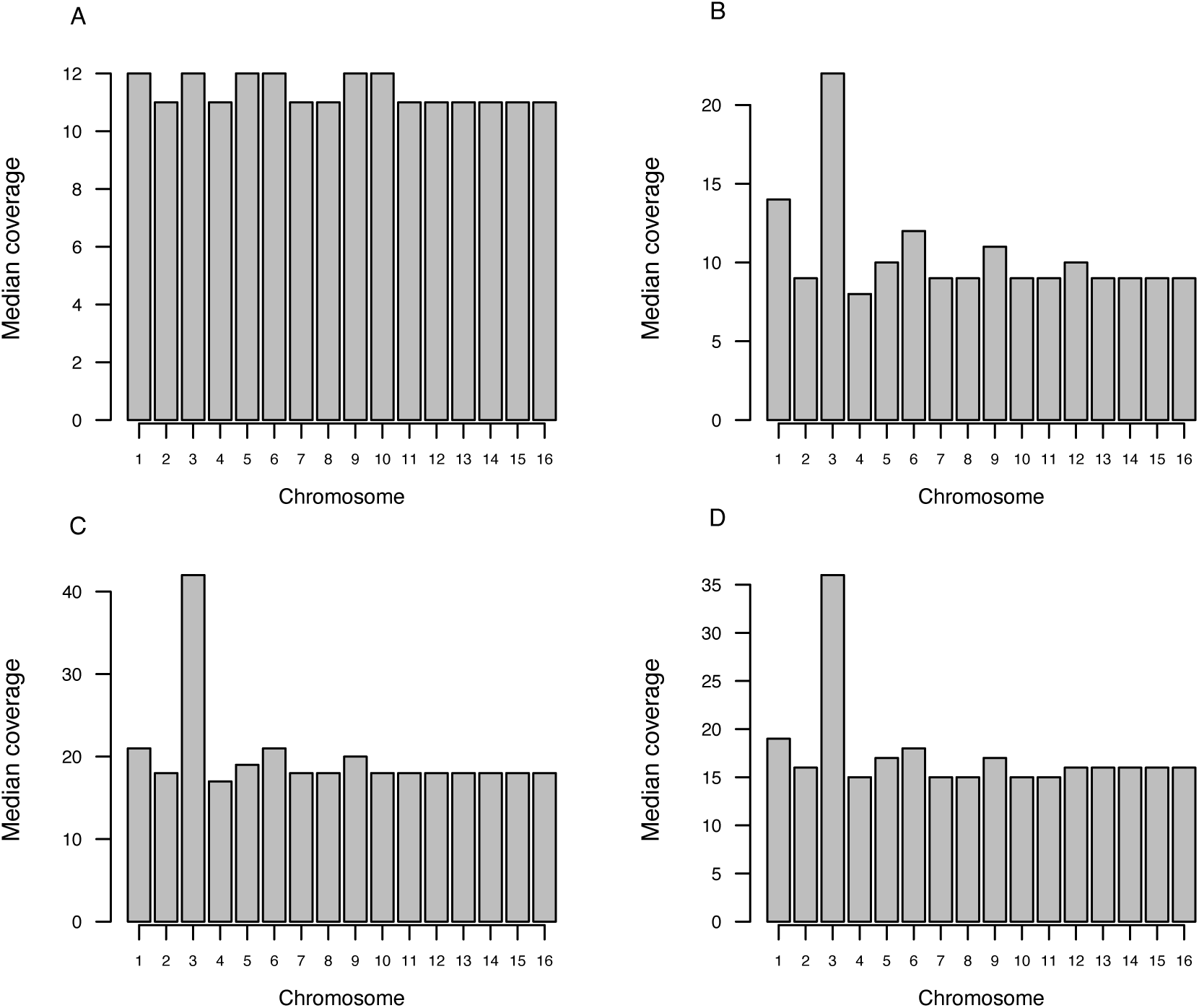
Coverage by chromosome indicates aneuploidy. Each bar represents the median sequencing coverage for a given chromosome. In the ancestral diploid (A), the chromosomes all have similar coverage. In the three populations from the sefling treatment that were sequenced (B–D), coverage on chromosome 3 is elevated by approximately 2-fold. Given flow cytometry results showing near-haploid genome content in the selfing populations, these results indicate disomy for chromosome 3.

**Figure S5:**
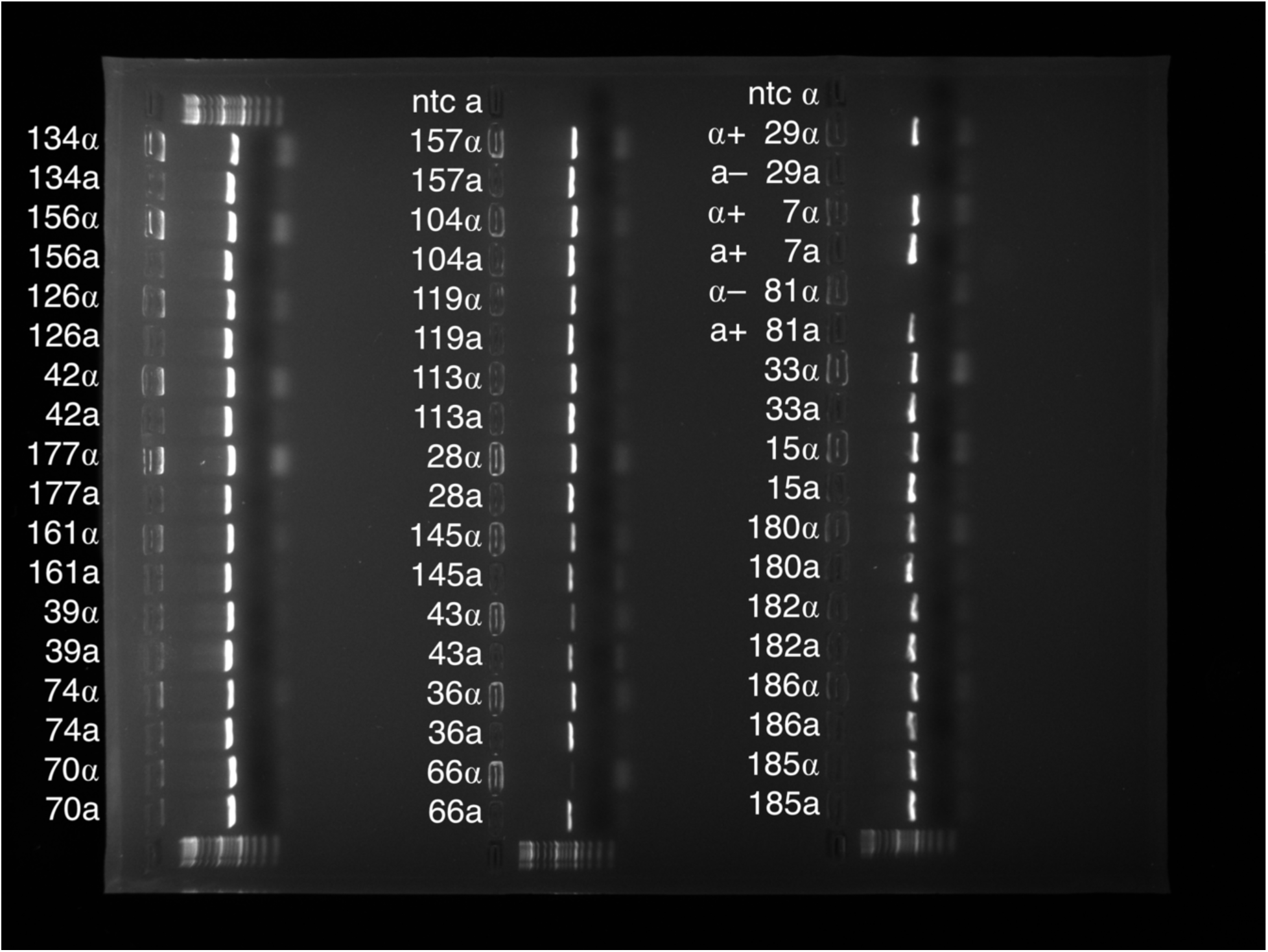
PCR of *MAT* in selfing lines. In each row, the strain ID is followed by the abbreviation a or ⍺ depending on which primer was used. “ntc” stands for no template control. 43⍺ and 66⍺ have faint but existing bands. Strain 29 acts as a positive MAT⍺ and negative MATa control. Strain 7 acts as a positive control for both mating type alleles, and strain 81 is our negative MAT⍺ and positive MATa control.

**Figure S6.**
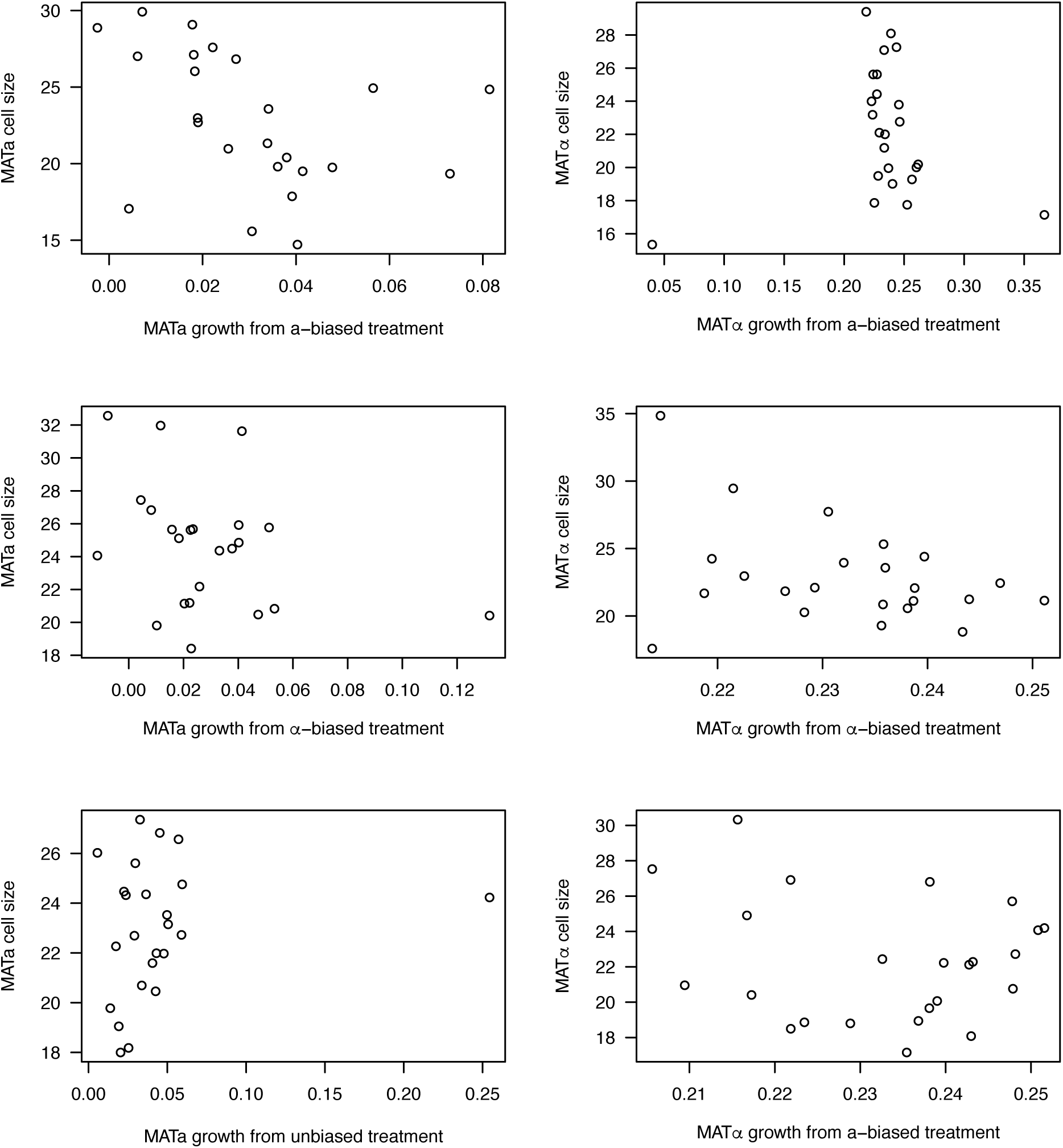
Correlations between cell size and growth rate. We found a significant negative average rank correlation between cell size and growth rate in the outcrossing treatments, driven by the biased mating ratio treatments, which showed a more negative correlation than the unbiased treatment. Faster growth was associated with smaller cell size among the populations that evolved under biased mating ratios.

**Figure S7.**
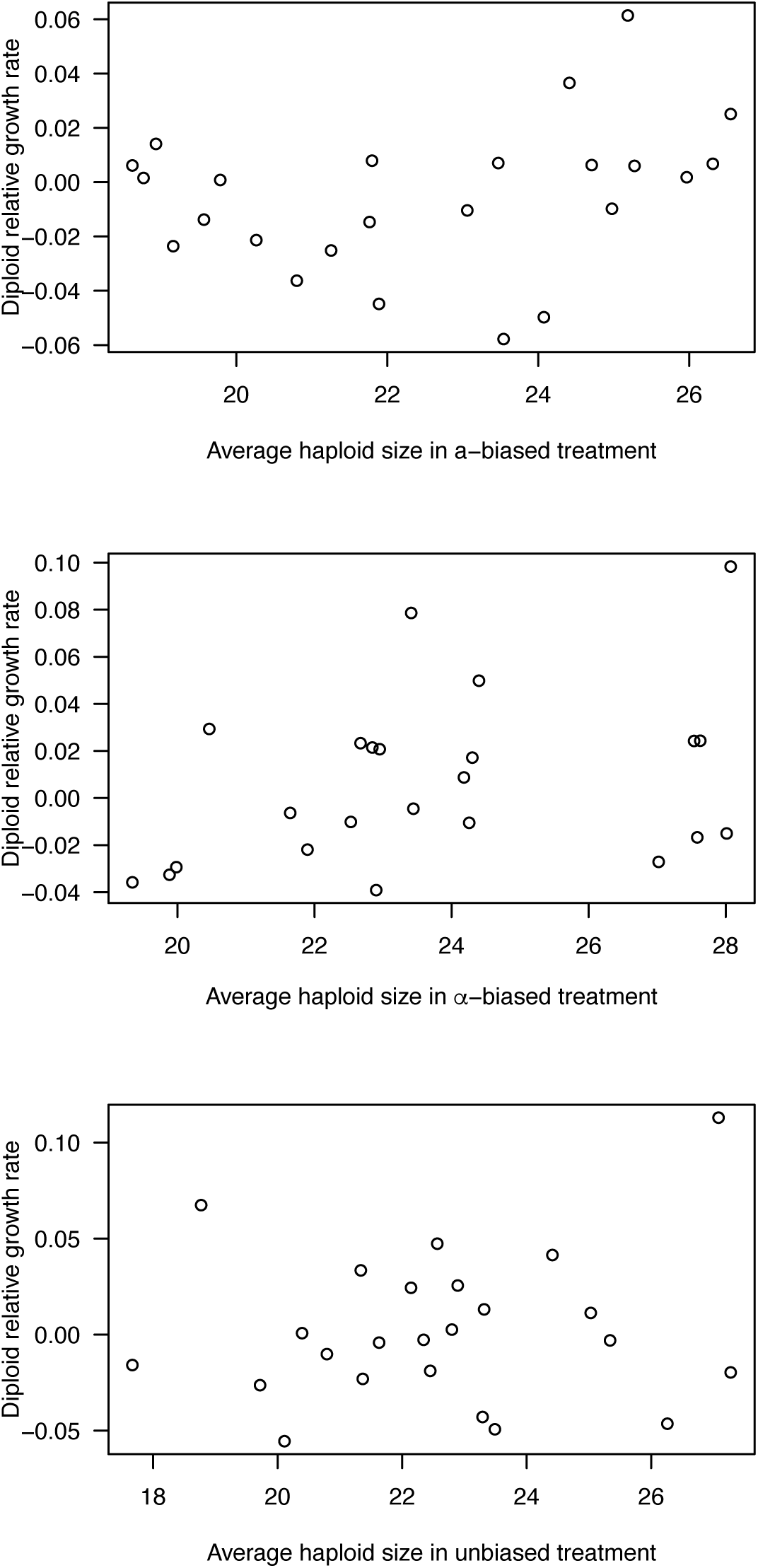
Correlations between diploid growth rate and haploid cell size. The growth rate of diploids was positively correlated with the average cell size of the constituent haploids (significant positive average rank correlation).

**Figure S8.**
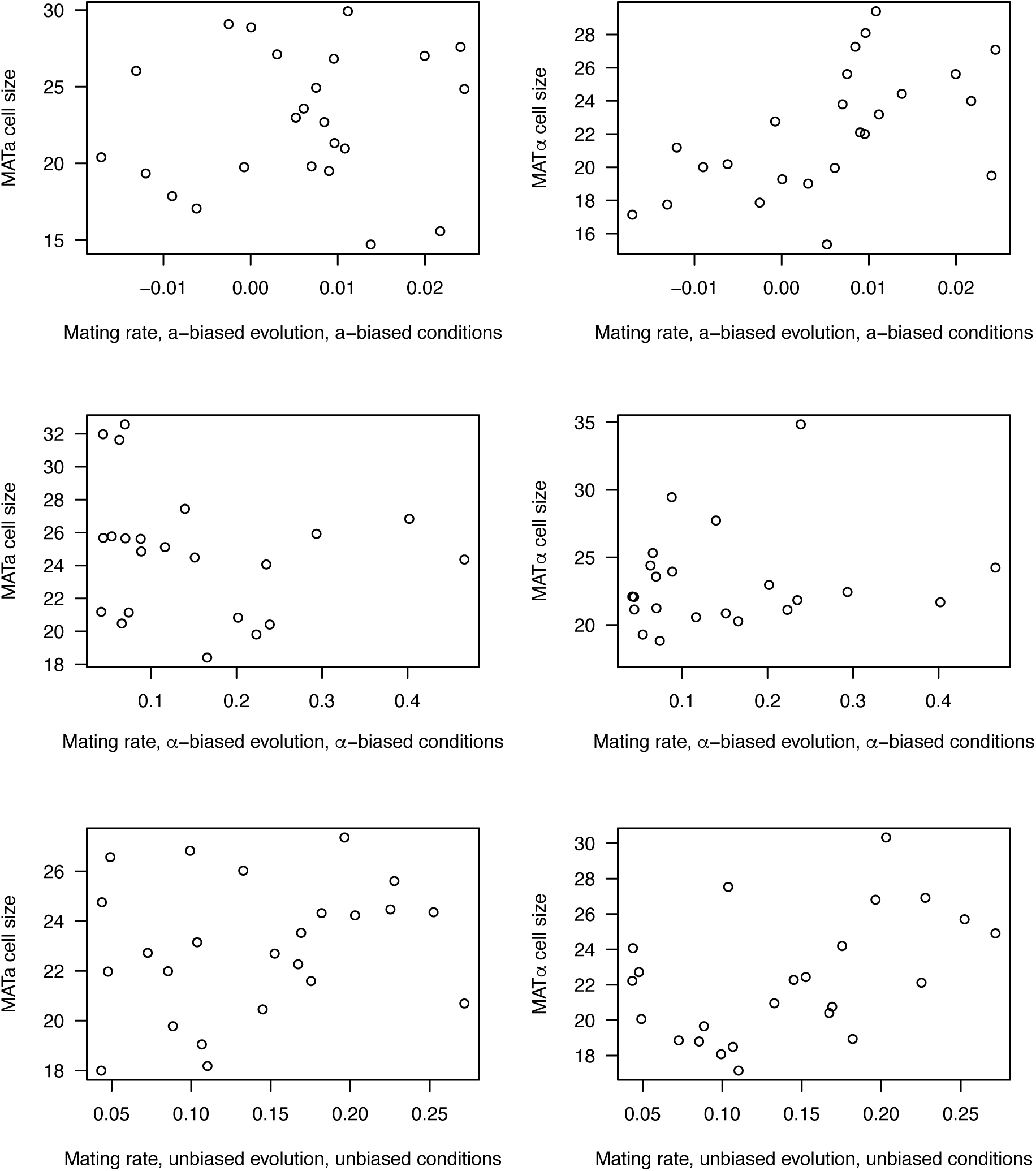
Correlations between cell size and mating rate. Cell size was positively correlated with mating rate, tested under the evolved mating-type ratio assay conditions, averaging rank correlations across treatments and mating types.

**Figure S9.**
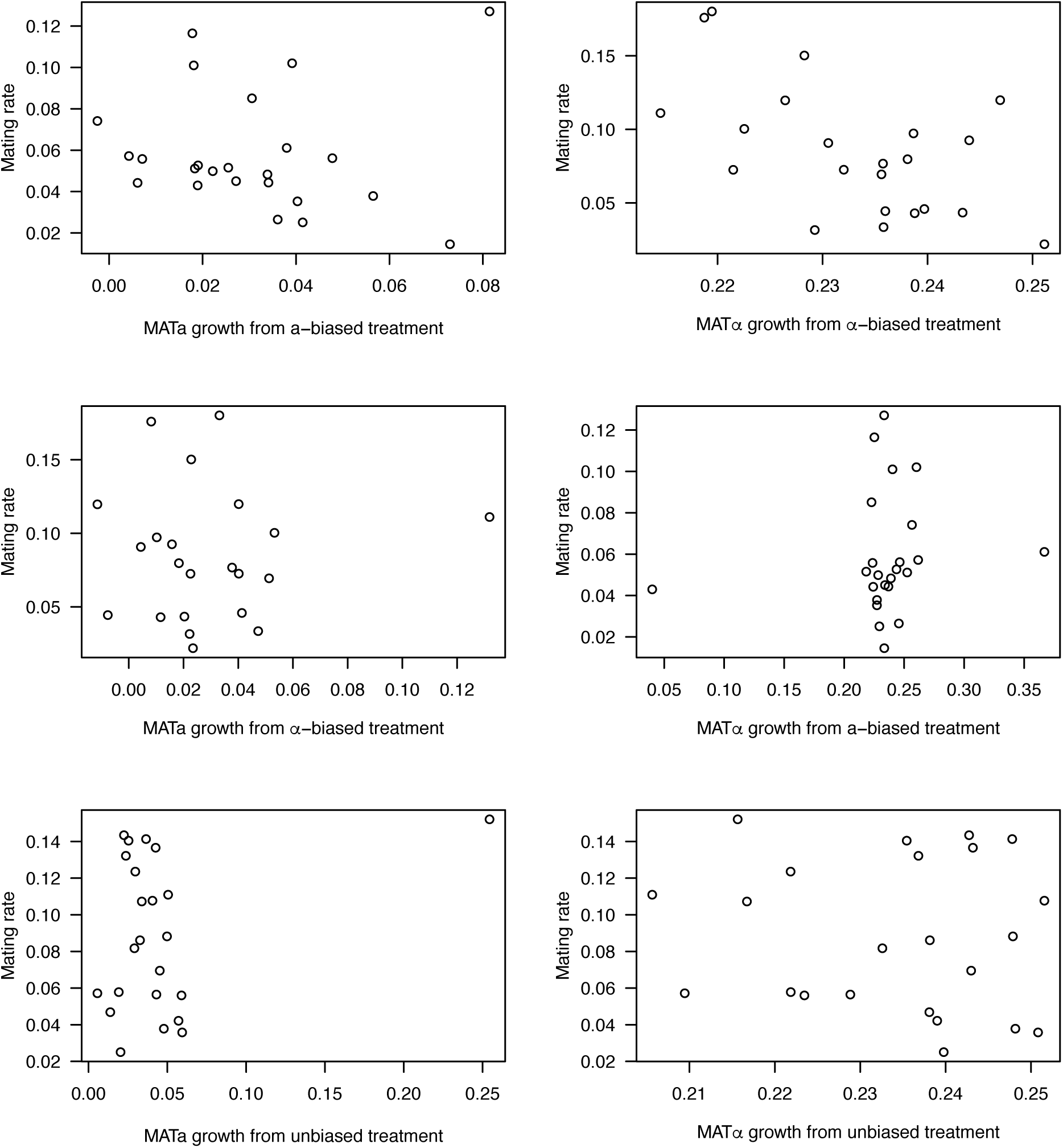
Correlations between mating rate and haploid growth rate. We found a significant negative average rank correlation between growth rate of the common cell type in selective media and the average mating rate across assay conditions, whereas there was no significant correlation for the rare cell type. In populations from the unbiased outcrossing treatment, we did not detect any significant correlation between growth and mating.

**Figure S10.**
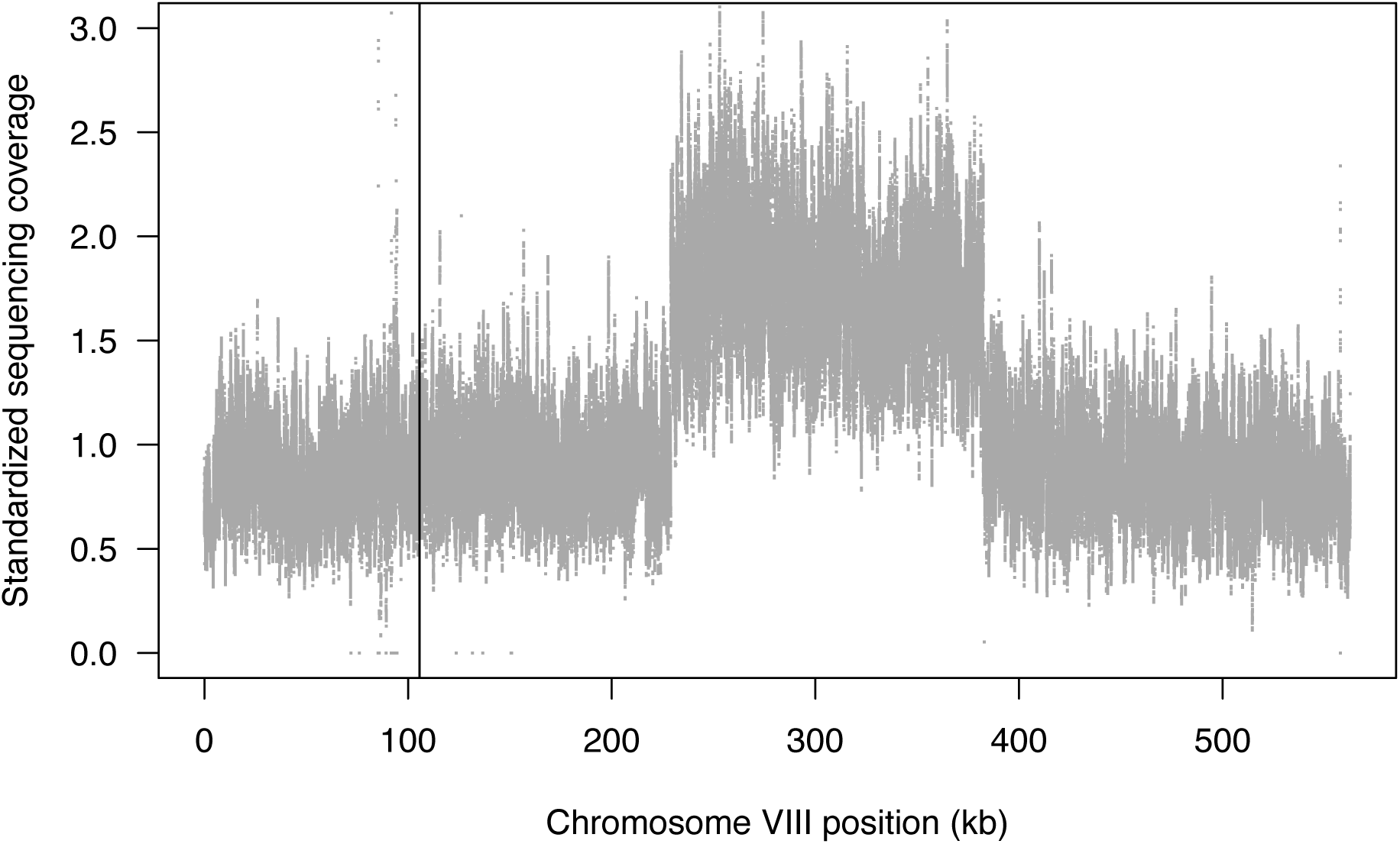
Evidence for a large duplication. The plot shows relative sequencing coverage at each site on chromosome VIII in a haploid sample, standardized to the average across samples. Coverage is elevated by approximately two-fold in the region 228965–382489, indicating that this region experienced a duplication.

**Table S1.** Experimental design. The colors in the table designate the type of medium used: permissive SE (green), diploid-specific (pink), presporulation (light blue), sporulation (purple), mating type-specific (yellow). All transfers were performed by placing 10 µL of culture in 1 mL of fresh medium. The colours in the table designate the type of medium used: permissive SE (green), diploid-specific (pink), presporulation (light blue), sporulation (purple), mating type-specific (yellow). All transfers were done by placing 10 µL of culture in 1mL of fresh medium.

| Day | Evolutionary treatment |  |
| --- | --- | --- |
|  | Delayed mating | Immediate mating |
| 1 | Growth (SE) |  |
| 2 am<sup>a</sup> | Mating | Growth (SE) |
| pm | Diploid selection |  |
| 3 am |  |  |
| pm<sup>a</sup> | Presporulation |  |
| 4 | Sporulation |  |
| 5 |  |  |
| 6 am | Ascus digestion |  |
| pm | Germination (MAT-specific) | Germination (SE) |
| 7 | MAT-specific selection | Diploid selection |

<sup>a</sup> Before mating and before transfer to presporulation medium, 200 μL of each culture was sampled for storage and freezing in 15% glycerol.

